# A photorespiratory glyoxylate shunt in the cytosol supports photosynthesis and plant growth under high light conditions in Arabidopsis

**DOI:** 10.1101/2024.12.03.626664

**Authors:** Xiaotong Jiang, Amanda M. Koenig, Berkley J. Walker, Jianping Hu

## Abstract

Photorespiration, a central aspect of plant metabolism that is tightly connected to photosynthesis, functions in part to support photosynthetic performance, especially under stress conditions such as high light. However, our understanding of the mechanisms underlying the role and regulation of photorespiration in plant response to high light is limited. To identify modulators of photorespiration under high light, we isolated genetic suppressors of the photorespiratory mutant *hpr1*, which is defective in the peroxisomal hydroxypyruvate reductase 1. A suppressor that partially rescued *hpr1* under high light was mapped to *GLYR1*, which encodes the cytosolic glyoxylate reductase 1 enzyme that converts glyoxylate to glycolate. Independent *GLYR1* loss-of-function mutants also partially rescued *hpr1* and another photorespiratory mutant, *catalase 2*. Our genetic, transcriptomic and metabolic profiling analyses together suggested a novel connection between cytosolic glyoxylate and a non-canonical photorespiratory route mediated by the cytosolic HPR2 enzyme, which we named the photorespiratory glyoxylate shunt. This shunt is especially critical under high light intensities when a high rate of photorespiratory flux is required and in the absence of a properly functional major photorespiratory pathway. Our findings support the metabolic flexibility of photorespiration and may help future efforts to improve crop performance under stress.

## Introduction

Light is an essential energy source and a critical environmental cue for plants, but its intensity often fluctuates beyond the ranges optimal for plant growth. For example, plants frequently experience high light conditions that exceed 2000 μmol m^-2^ s^-1^ on sunny days, which pose significant stress to plants in the field^1,2^. When light intensities surpass the photosynthetic capacity, generation of reactive oxygen species (ROS) from the photosynthetic apparatus is enhanced, which can cause photodamage to protein complexes and subsequently inactivate the electron transport chain^3,4^. To cope with this stress, plants have developed strategies to respond at various levels, such as the antioxidant system to scavenge ROS, dissipation of excessive energy as non-photochemical quenching, cyclic electron flow to balance ATP/NADPH production, transcriptional reprogramming, chloroplast and leaf movement, and anthocyanin accumulation^5,6^.

Photorespiration, a metabolic process closely related to photosynthesis, is initiated after the oxygenation of ribulose 1,5-bisphosphate (RuBP) by the photosynthetic enzyme ribulose 1,5-bisphosphate carboxylase-oxygenase (rubisco)^7^. Through a series of reactions residing sequentially in the chloroplast, peroxisome, mitochondrion, and the cytosol, the oxygenation product 2-phosphoglycolate (2-PG), which inhibits cell functions, is eventually converted to 3-phosphoglycerate (3-PGA) to be recycled back to the Calvin-Benson cycle in the chloroplast^7^ (Fig. 1).

**Fig. 1.**
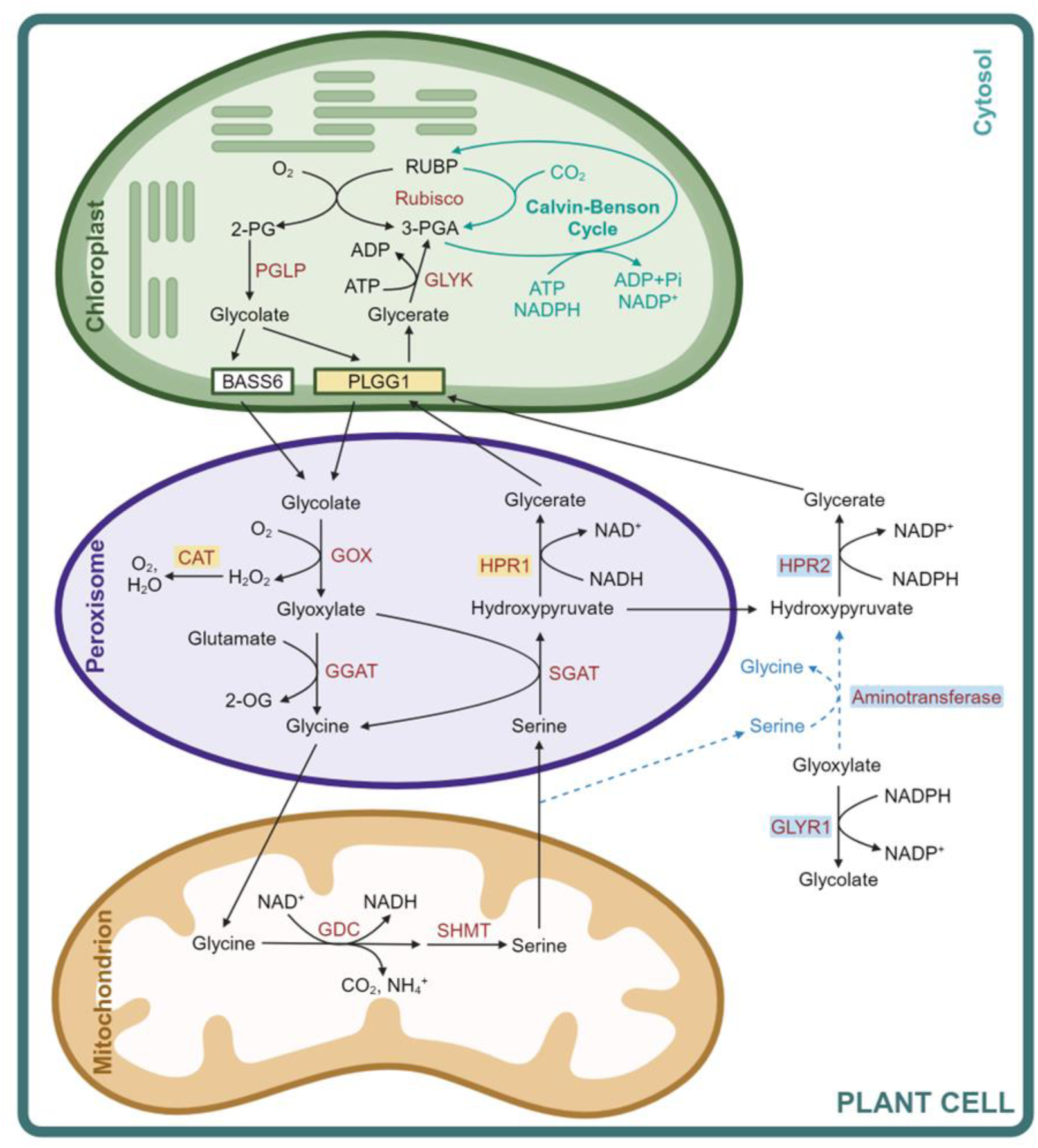
The established photorespiration pathway and the cytosolic glyoxylate shunt proposed in this study. Photorespiration involves a series of reactions in the chloroplast, peroxisome, mitochondrion, and cytosol. We propose that defective GLYR1 allows the accumulated free glyoxylate in the cytosol to react with serine, catalyzed by an unknown aminotransferase. The hydroxypyruvate produced can be further converted by HPR2 to glycerate, which re-enters the chloroplast. Abbreviations: 2-OG, 2-oxoglutarate; 2-PG, 2-phosphoglycolate; 3-PGA, 3-phosphoglycerate; BASS6, bile acid sodium symporter 6; CAT, catalase; GDC, glycine decarboxylase complex; GGAT, glutamate:glyoxylate aminotransferase; GLYK, glycerate kinase; GLYR1, glyoxylate reductase 1; GOX, glycolate oxidase; HPR, hydroxypyruvate reductase; PGLP, 2-PG phosphatase; PLGG1, plastidial glycolate/glycerate transporter 1; Rubisco, RuBP carboxylase/oxygenase; RuBP, ribulose-1,5-bisphosphate; SGAT, serine:glyoxylate aminotransferase; SHMT, serine hydroxymethyltransferase. Created with BioRender.com.

Although photorespiration is often considered a sub-optimal process because it consumes energy and releases pre-fixed carbon as CO_2_, a properly functional photorespiratory pathway supports photosynthetic performance, especially under stress conditions. In ambient air and under regular growth conditions, many photorespiratory mutants exhibit compromised photosynthesis and growth, phenotypes that are largely recovered under high CO_2_ environments where rubisco oxygenation is inhibited^8,9^. Although the exact reasons for these air-grown phenotypes are unclear, the accumulated photorespiratory intermediates in these mutants, such as 2-PG, glyoxylate, and glycerate, can inhibit the activities of photosynthesis-related enzymes^9^. Additionally, several Arabidopsis photorespiratory mutants, including *hydroxypyruvate reductase 1* (*hpr1*), *plastidial glycolate/glycerate transporter 1* (*plgg1*), *catalase 2* (*cat2*) and *glycolate oxidase 1* (*gox1*), have much more severe photosynthetic phenotypes under high and dynamic light compared with low and constant light, supporting the increased importance of a properly functional photorespiratory pathway under high light^10^. Under stress, photorespiration is believed to function as an alternative electron sink that consumes excessive energy produced by the photosynthetic light reactions^11–14^, although a recent work challenged the idea that photorespiration’s role as an alternative electron sink is photoprotective^15^.

As a core enzyme in photorespiration, HPR catalyzes the reduction of hydroxypyruvate to produce glycerate (Fig. 1). In Arabidopsis, three members of the HPR family have been shown to function in photorespiration^16,17^. Peroxisomal HPR1 plays a major role in reducing hydroxypyruvate with NADH as the co-factor, whereas HPR2 and HPR3 in the cytosol have low activities and higher affinities for NADPH^16–19^. HPR enzymes are important in photosynthesis, because the knockout mutant of *HPR1* shows compromised photosynthetic performance and growth in ambient air, with additive effects observed in the *hpr1 hpr2* double and *hpr1 hpr2 hpr3* triple mutants^16,17^. Under high light, *hpr1* exhibits stronger phenotypes, including decreases in the efficiency of photosystem II (PSII), increases in non-photochemical quenching, activation of cyclic electron flow, accumulation of H_2_O_2_, and a marked reduction in chlorophyll and anthocyanin^10,19^. Further investigation of *hpr1* under high light found that 2-PG accumulation inhibits the activity of triose phosphate isomerase, an enzyme of the Calvin-Benson cycle that converts glyceraldehyde 3-phosphate to dihydroxyacetone phosphate. This inhibition results in a cytosolic bypass and glucose-6-phosphate (G6P) shunt in the Calvin-Benson cycle where the G6P shunt consumes ATP, triggering high rates of cyclic electron flow to balance energy demand^20^. Consistent with the activation of the G6P shunt, increased CO_2_ release was also observed in *hpr1*^21,22^. However, there are other possible explanations for this extra CO_2_ release, as the non-enzymatic decarboxylation of hydroxypyruvate and serine consumption through serine decarboxylase also seem to occur in *hpr1*^21,22^. Finally, HPR1 was found to maintain the repair of PSII under high light^19^.

HPR enzymes have a broad influence on the level of metabolites in the plant. Deficiency in one or more HPRs increases the level of most photorespiratory intermediates, including glycolate, glycine, serine, hydroxypyruvate and glycerate, and these metabolic phenotypes are affected by photoperiods^16,17,22^. Consistent with the impaired photosynthesis, carbohydrate levels are largely decreased in the *hpr1* mutant^22^. Levels for other metabolites related to photorespiration, such as intermediates in the tricarboxylic acid cycle and many amino acids, are elevated or reduced in *hpr1*^16,17,22^. Interestingly, different from its daytime-dependent accumulation pattern in the wild type plants, serine was found to be constitutively elevated in the *hpr1* mutant, inhibiting the expression of photorespiratory genes and reducing the level of the corresponding enzymes^23^. Additionally, the accumulated glycolate can replace the bicarbonate ligand in PSII in *hpr1*, shifting the midpoint potential of the quinone acceptor and reducing the generation of single oxygen^24^.

While the main flux of photorespiration is well known, there are examples of alternative fluxes, such as the aforementioned cytosolic HPR2 and HPR3 enzymes that are partially redundant in function with HPR1. Understanding other routes of carbon flux associated with photorespiration is important to fully deciphering this metabolic network. In addition, photorespiration is tightly linked to primary metabolic pathways and crucial to plant survival under certain environmental conditions, thus this pathway is expected to be regulated. Although research in this area is still scarce, current evidence indicates that photorespiration responds to environmental factors such as CO_2_, O_2_ and light, receives feedback from its own metabolites and enzymes, and is regulated at transcriptional, post-transcriptional and post-translational levels^25–27^. Taken together, mechanistic research into the flexibility and regulation of photorespiration will be highly valuable to completely elucidating the role of this pathway in plant physiology and plant interaction with the environment.

In this work, we investigated the regulation and flexibility of photorespiration by characterizing a genetic suppressor of the Arabidopsis *hpr1* mutant under high light, mapping the underlying gene, and conducting follow-up genetic, metabolic flux and transcriptome analyses. We found that defective glyoxylate reductase 1 (GLYR1), a cytosolic enzyme that catalyzes the conversion of glyoxylate to glycolate, can partially rescue the mutant phenotypes of *hpr1* in plant growth, photosynthesis, levels of photorespiratory metabolites, and transcriptome under high light conditions. Further examination showed that deficient *GLYR1* can also partially suppress the phenotype of the catalase mutant *cat2*, but not that of the glycolate/glycerate transporter mutant *plgg1*. Combining transitional metabolic profiling, glyoxylate feeding, and genetic analyses, we provided evidence for a cytosolic photorespiratory shunt that converts accumulated glyoxylate to hydroxypyruvate, which can act, at least partially, through the cytosolic HPR2 enzyme to enhance carbon recycling. This cytosolic shunt seems to be especially critical under high light intensities, when a high rate of photorespiratory flux is required to deal with increased rates of total rubisco oxygenation reaction, and in the absence of a functional major photorespiratory pathway. Our findings suggest that the metabolic flexibility of photorespiration can help plants adjust to stress conditions, thus may assist in future efforts to improve plant performance and stress tolerance.

## Results

### Loss of/reduced function of glyoxylate reductase 1 (GLYR1) partially rescues the growth and metabolic phenotypes of the *hpr1* mutant

To identify proteins with a regulatory or modulatory role in photorespiration, we performed EMS mutagenesis on Arabidopsis *hpr1-1* mutant seeds and screened for genetic suppressors based on their growth and photosynthetic phenotypes under high light conditions (∼700 μmol m^-2^ s^-1^). One suppressor, *shpr7* (*hpr1 suppressor number 7*), was found to partially suppress the small rosette size of *hpr1-1* under high light relative to normal light conditions (100 μmol m^-2^ s^-1^) (Fig. 2a and b).

**Fig. 2.**
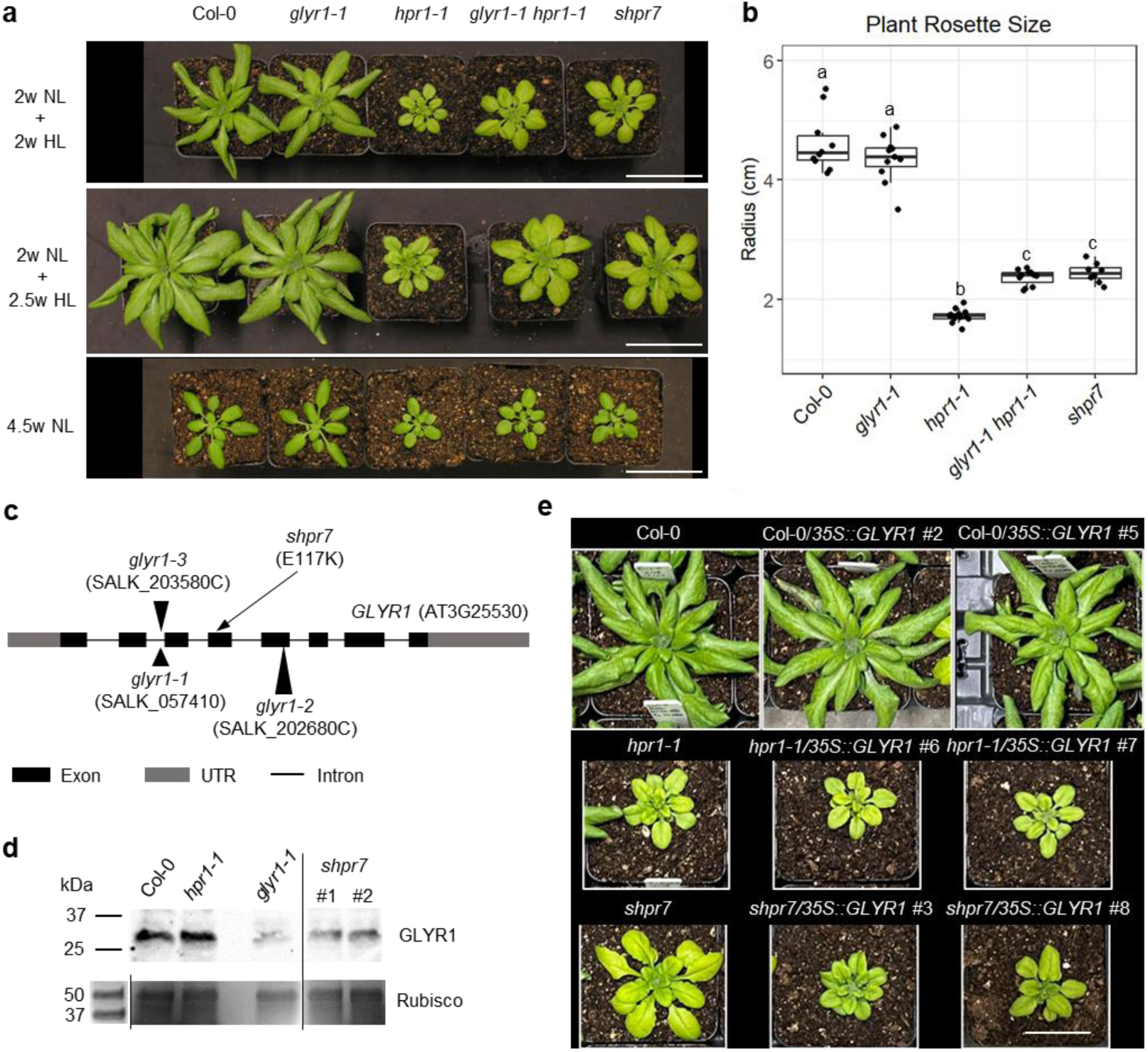
Defective GLYR1 partially rescues the growth phenotypes of *hpr1*. **(a)** Plants grown for 2 weeks (2w) under normal light (NL, 100 µmol m^-2^ s^-1^) followed by 2 or 2.5 weeks of growth under high light (HL, 700 µmol m^-2^s^-1^), or constantly grown under normal light for 4.5 weeks. Scale bars = 5 cm. **(b)** Radius measurements of the total rosette of 4-week-old plants grown under 2 weeks of normal light followed by 2 weeks of high light. Different letters indicate statistically significant differences (p<0.05), which were determined by One-way ANOVA with Tukey’s HSD test. Biological replicates: n=10 for Col-0, n=11 for *glyr1-1* and *glyr1-1 hpr1-1*, n=13 for *hpr1-1*, and n=8 for *shpr7*. **(c)** Schematic depiction of the *GLYR1* gene and positions of the mutations in various alleles. **(d)** Immunoblot analysis of the GLYR1 protein from different genotypes. The GLYR1 protein was detected by an anti-GLYR1 peptide antibody (top). Rubisco stained by Coomassie Blue in the SDS-PAGE gel was used as the loading control (bottom). **(e)** Overexpressing *GLYR1* reverts the suppression phenotype back to the *hpr1* mutant phenotype. Plants were grown under 2 weeks of normal light followed by 2.5 weeks of high light. Scale bars = 3 cm.

To characterize the causal mutation in *shpr7*, we backcrossed *shpr7* to *hpr1-1*. The segregation ratio of *hpr1*-like vs. suppressor-like plants in the BC_1_F_2_ generation was 3:1, suggesting that the suppression is caused by a recessive mutation. To map the gene responsible for the suppression, genomic DNA extracted from 80 suppressor-like individuals in the BC_3_F_2_ generation was pooled for whole-genome sequencing. We identified a point mutation in the exon of the *Glyoxylate Reductase 1* (*GLYR1*) gene, which encodes an NADPH-dependent glyoxylate/succinic semialdehyde reductase ^28,29^, causing a glutamate (E)-to-lysine (K) substitution at amino acid 117 (Fig. 2c). Reverse transcription PCR (RT-PCR) analysis detected similar levels of the *GLYR1* transcripts in *shpr7* and the wild type (Supplementary Fig. 1a), suggesting that this mutation does not lead to significant changes in *GLYR1* gene expression. To determine the effect of the E117K mutation at the protein level, peptide antibodies against amino acids 229-243 were generated, which detected an apparently decreased level of the GLYR1 protein (30.7kDa) in high light-grown *shpr7* (Fig. 2d), indicating that this point mutation may cause the protein to be less stable. The antigen peptide is from a highly identical region between AtGLYR1 and its homolog, AtGLYR2, where the two proteins differ only by one amino acid^30^. We therefore reasoned that the weak signal in *glyr1-1* in the immunoblot might be derived from the mature form of GLYR2, after its N-terminal transit peptide for chloroplast and mitochondrial dual targeting is removed upon import into the organelles (Fig. 2d).

To confirm that loss-of-function mutations in *GLYR1* can lead to the suppression of *hpr1-1* like that shown in *shpr7*, we obtained three independent T-DNA insertion mutant lines of *GLYR1* (Fig. 2c). All three lines lacked detectable full-length *GLYR1* transcript (Supplementary Fig. 1b) and showed comparable growth and morphologies to the wild type (Col-0) plants (Supplementary Fig. 1c). Double mutants generated by crossing these lines individually with *hpr1-1* all exhibited very similar phenotypes as *shpr7* (Fig. 2a and b, Supplementary Fig. 1c), confirming that loss of function of GLYR1 can partially rescue the *hpr1-1* mutant phenotypes. Overexpressing the *GLYR1* gene (*35S::GLYR1*) in *shpr7* reverted its rosette size back to *hpr1*-like under high light (Fig. 2e, Supplementary Fig. 1d), further confirming *GLYR1* as the causal gene for the suppression of *hpr1*. Since the three T-DNA mutants were indistinguishable from each other in their capability to suppress *hpr1-1,* we selected the *glyr1-1* allele for follow-up experiments.

GLYR1 was shown to catalyze the conversion of glyoxylate to glycolate, and succinic semialdehyde to γ-hydroxybutyrate^29^. Because glyoxylate and glycolate are photorespiratory intermediates, we tested if the deficiency in GLYR1 also causes metabolic changes in the photorespiratory pathway. The *hpr1-1* mutant grown under high light conditions had elevated levels of all the 6 photorespiratory metabolites tested (Fig. 3), consistent with previous reports that knockout mutants of *HPR1* accumulate photorespiratory intermediates under normal growth conditions^16,22^. The suppressor *shpr7* and the double mutant *glyr1-1 hpr1-1* showed partial rescue of the accumulation of at least 5 of these metabolites, including glycolate, glyoxylate, serine, hydroxypyruvate, and glycerate, and consistent with its wild type-like plant appearance, the *glyr1-1* single mutant contained similar levels of these metabolites as Col-0 (Fig. 3).

**Fig. 3.**
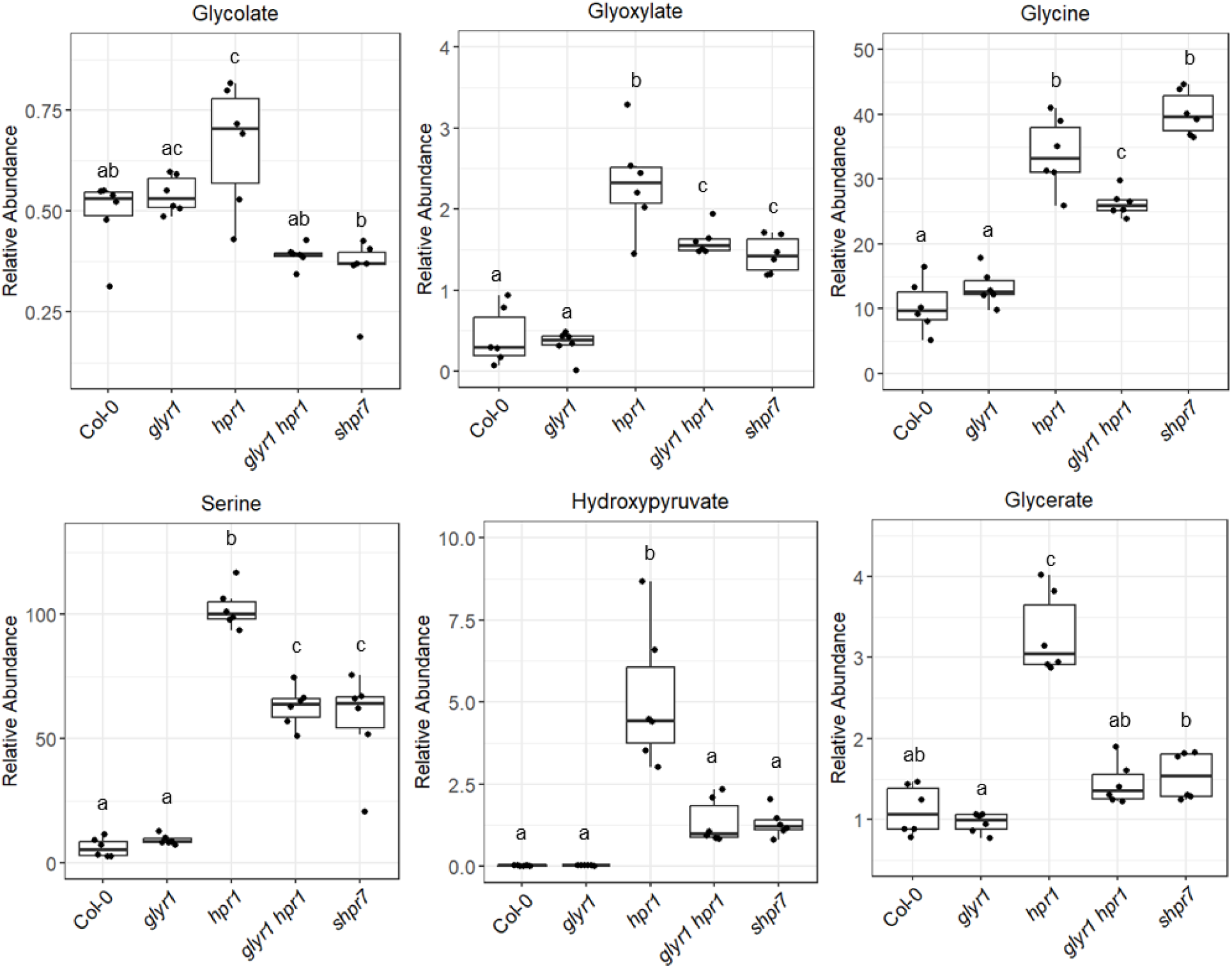
Profiling of stable photorespiratory metabolites in 4-week-old high light-treated plants. Abnormal levels of the photorespiratory intermediates in *hpr1* are mostly partially rescued by deficient GLYR1. Plants were grown under 2-week normal light followed by 2-week high light. Different letters indicate statistically significant differences (p<0.05), which were determined by One-way ANOVA with Tukey’s HSD test. Biological replicates: n=6.

Taken together, our results demonstrated that deficiencies in the Arabidopsis GLYR1 protein can partially suppress the mutant phenotypes of *hpr1* in both plant growth and levels of the photorespiratory metabolites, especially under high light conditions.

### Deficient GLYR1 largely reverts the transcriptional reprogramming of the *hpr1* mutant under short-term photorespiratory conditions

The suppression of *hpr1*’s growth and metabolic phenotypes by deficient GLYR1 prompted us to further investigate the global impact of *glyr1* on the Arabidopsis transcriptome. We performed RNA-seq to analyze the transcriptomes of Col-0, *hpr1-1*, *glyr1-1*, *glyr1-1 hpr1-1,* and *shpr7* under short-term photorespiratory conditions, as prolonged stress conditions may result in secondary effects that obscure the direct impact of the mutations during photorespiration. To this end, plants were grown for 3 weeks under normal light and high CO_2_ (2,000 PPM CO_2_), where photorespiration is largely inhibited (Supplementary Fig. 2). At the end of the dark period, plants were transferred to ambient air and high light conditions to induce photorespiration. Leaf samples harvested after 3 h and 10 h of the treatment were used for RNA-seq.

We identified 10,139 and 16,104 differentially expressed genes (DEGs), at 3 h and 10 h respectively, between *hpr1-1* and Col-0 (Fig. 4a and b), constituting approximately 40%-60% of the ∼25,500 Arabidopsis genes. This large number of DEGs demonstrates the remarkable transcriptional reprogramming in *hpr1-1*. Compared to Col-0, the *glyr1-1* single mutant had minimal DEGs, i.e., 59 at 3 h and 12 at 10 h. However, the same *glyr1-1* mutation dramatically altered gene expression in the *hpr1-1* background. For example, at 10 h, over half of the genes in Arabidopsis are differentially expressed in *glyr1-1 hpr1-1* (15,955) vs. *hpr1-1* as well as in *shpr7* (15,795) compared to *hpr1-1*, highlighting the strong impact of deficient GLYR1 in rescuing *hpr1-1* at the transcriptional level.

**Fig. 4.**
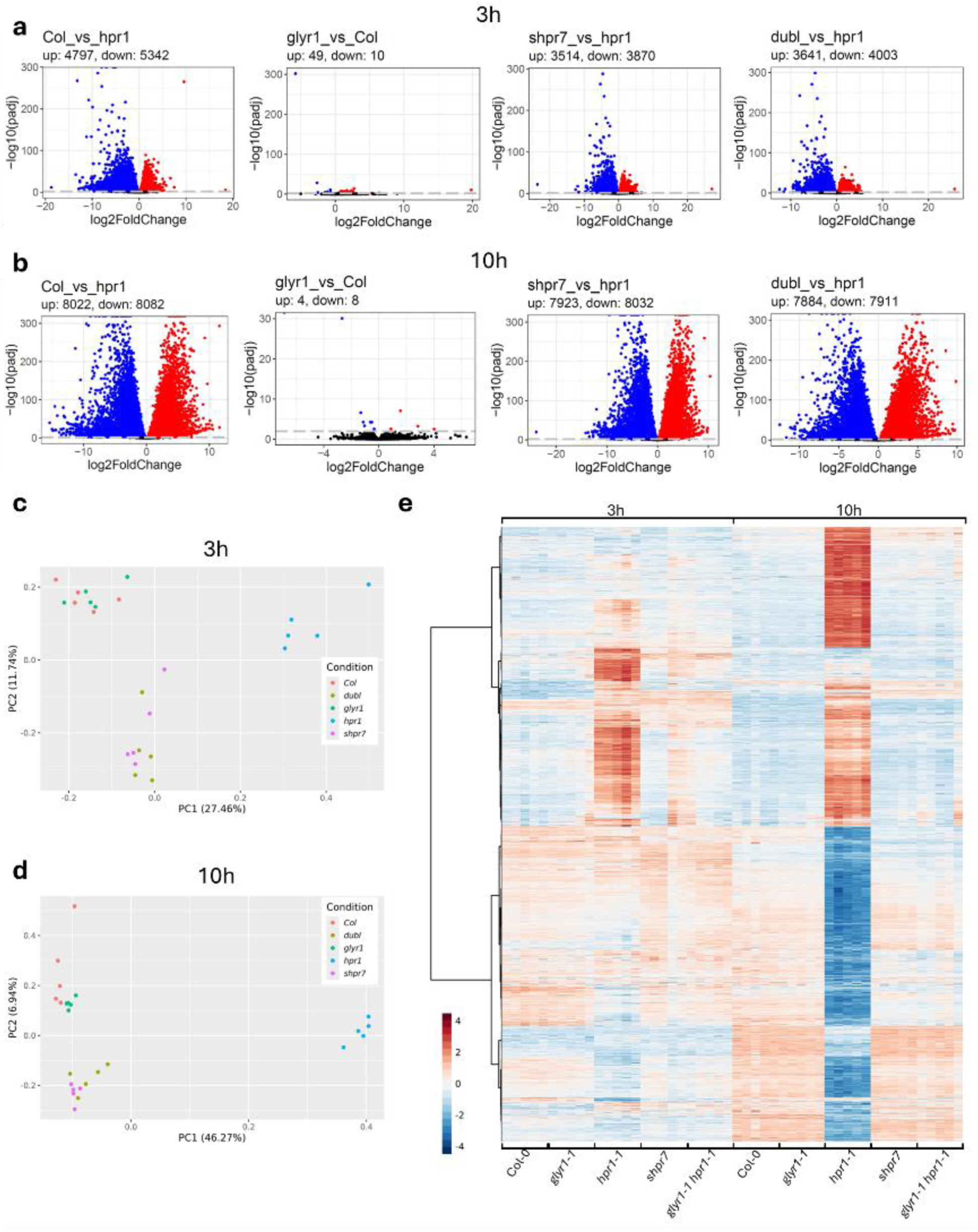
Deficient GLYR1 largely reverts the transcriptional reprogramming in *hpr1*. Plants were grown for 3 weeks under high CO_2_ in normal light, and then transferred to ambient air and high light during the dark period. Leaf tissue was sampled after 3 h and 10 h for RNA-seq. Biological replicates: n=5. **(a-b)** Volcano plots displaying differentially expressed genes (DEGs) in four pairwise comparisons: *glyr1-1* vs. Col-0, and Col-0, *shpr7* and *glyr1-1 hpr1-1* vs. *hpr1-1* at 3 h **(a)** and 10 h **(b)**. Genes that have an absolute log2 fold ratio>0 and an adjusted *p-value* (padj) of < 0.01 (represented logarithmically on the y-axis) are considered DEGs and shown above the gray dashed line. Blue, red, and black dots represent the down-regulated, up-regulated, and the unchanged genes, respectively. **(c-d)** Principal Component Analysis (PCA) at 3 h **(c)** and 10 h **(d).** Col-0 and *glyr1-1* cluster together and *shpr7* and *glyr1-1 hpr1-1* (dubl) cluster together, with both groups clustering discretely from *hpr1-1.* **(e)** A heatmap in which *hpr1-1* shows dramatic transcriptional reprogramming that is restored to near Col-0 levels in *glyr1-1 hpr1-1* (dubl) and *shpr7*. Genes with at least a 4-fold change in the Col-0 vs. *hpr1-1* comparison were selected and their expression levels across time points and genotypes are represented.

Principal component analysis (PCA) resolved the samples into discrete clusters according to their predominating similarities and differences. At both timepoints, Col-0 and *glyr1-1*, and *glyr1-1 hpr1-1* and *shpr7*, were respectively clustered together, with both clusters clearly separated from *hpr1-1* (Fig. 4c and d), consistent with the growth phenotypes of these lines. The *hpr1-1* replicates segregated from the other genotypes according to the first principal component (PC1), which accounts for the most variation in the data (Fig. 4c and d), emphasizing that the transcriptomes of *glyr1-1 hpr1-1* and *shpr7* are discernably more similar to Col-0 than to *hpr1-1*.

Consistent with the clustering patterns in the PCA, the sweeping transcriptional reprogramming in *glyr1-1 hpr1-1* and *shpr7* recovers much of the transcriptome of *hpr1-1* to wild type expression levels during short-term photorespiratory conditions (Fig. 4e). At 3 h, more than 50% of the DEGs changed by at least 2-fold in *hpr1* (p_adj_ < 0.01 and |Log2Fold-Change| > 1) are already restored to Col-0 expression patterns by deficient GLYR1 (Supplementary Fig. 3a). Of the 12,230 DEGs with at least a 2-fold change in Col-0 compared to *hpr1-1* at 10 h, 11,373 genes are similarly up- or down-regulated in either *glyr1-1 hpr1-1* or *shpr7*, and 10,732 genes show the same differential expression patterns (up-/down-compared to *hpr1-1*) in all three genotypes (Supplementary. Fig. 3b). Among the overall expression patterns of DEGs with at least a 4-fold change in the Col-0 vs. *hpr1-1* comparison, it is immediately apparent that the *hpr1-1* mutant has a distinct expression profile compared to the other genotypes, while Col-0, *glyr1-1 hpr1-1*, and *shpr7* are strikingly similar (Fig. 4e). At 10 h, *hpr1-1* gene expression patterns are generally oppositely regulated compared to Col-0 and *glyr1-1*. Moreover, the gene expression profiles of *glyr1-1 hpr1-1* and *shpr7* almost entirely revert to Col-0 expression levels, indicating that the lack of functional GLYR1 suppresses the transcriptional changes observed in *hpr1-1* (Fig. 4e), including expression profiles of genes associated with photorespiration and photosynthesis (Supplementary Fig. 4).

Overall, our RNA-seq data revealed that *hpr1-1* causes massive transcriptional reprogramming under high light, and deficient GLYR1 globally rescues *hpr1-1* at the transcriptome level, including transcriptional changes in genes related to photorespiration and photosynthesis and many other biological functions.

### Deficient GLYR1 also partially rescues the phenotypes of the photorespiratory mutant *cat2*, but not those of *plgg1*

To investigate if the suppression of *hpr1* by *glyr1* is specifically linked to HPR1 or broadly connected to components of the photorespiratory pathway, we crossed *hpr1-1* into two other photorespiratory mutants, *cat2-1* and *plgg1-1,* which are defective in the peroxisomal catalase 2 enzyme and the chloroplast glycolate/glycerate transporter, respectively (Fig. 1). Similar to *hpr1-1*, *cat2-1* and *plgg1-1* are also compromised in growth and photosynthesis under high light^10^. Interestingly, under high light, the double mutant *glyr1-1 cat2-1* exhibited a bigger rosette size than *cat2-1*, whereas *glyr1-1* was unable to improve the growth of *plgg1-1* (Fig. 5a).

**Fig. 5.**
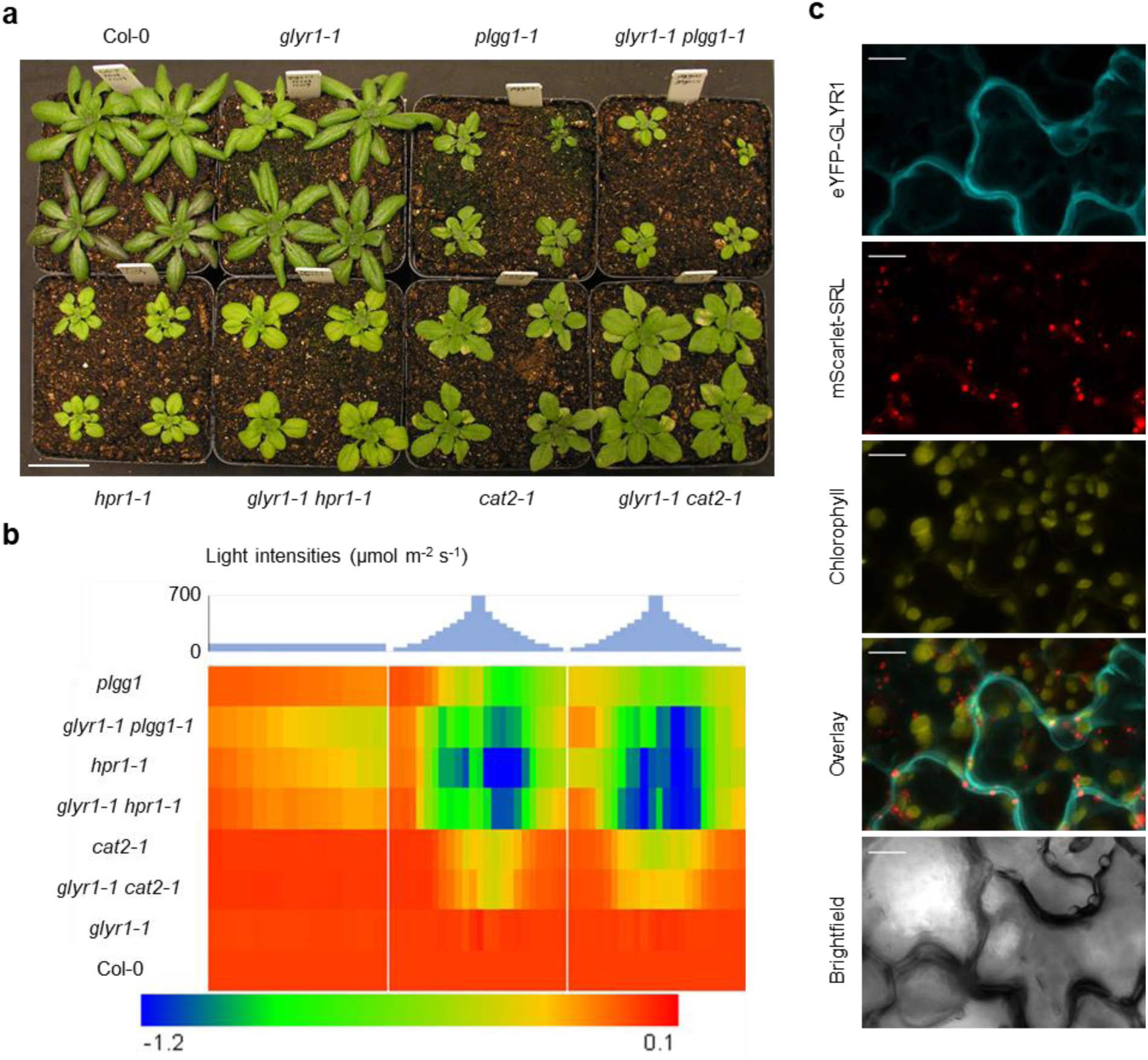
Analysis of the impact of defective GLYR1 on other photorespiratory mutants and GLYR1 protein localization. **(a)** Plants grown under 2 weeks of normal light followed by 2 weeks of high light. Scale bars = 3 cm. **(b)** Heatmap of the quantum efficiency of photosystem II for 2-week-old plants grown under dynamic and high light conditions. Values for the mutants were normalized to that of the wild-type Col-0. Biological replicates: n=12 for Col-0, *cat2-1*, *glyr1-1 cat2-1*, *glyr1-1 hpr1-1*, and *plgg1-1*; n=11 for *glyr1-1* and *hpr1-1*; and n=13 for *plgg1-1*. **(c)** Maximum intensity Z-projection of confocal images spanning 30 µm of the tobacco leaf tissue co-expressing eYFP**-**GLYR1 (cyan) and mScarlet-I-SRL (peroxisome marker, red). Chloroplast signals are from chlorophyll autofluorescence (yellow). Scale bar = 10 μm.

To determine if *glyr1* can also rescue the reduced photosynthetic efficiency in *cat2-1* and *plgg1-1*, as well as *hpr1-1*, we measured quantum efficiency of photosystem II (Φ_II_), a critical parameter of photosynthesis, in mutant and Col-0 seedlings. A 3-day light regime, with normal light on the first day and light gradients on days 2 and 3, was applied (Fig. 5b). Consistent with their growth phenotypes, *hpr1-1*, *cat2-1* and *plgg1-1* had lower Φ_II_ values, especially under higher light intensities, whereas *glyr1-1* resembled Col-0 (Fig. 5b). As expected, GLYR1 deficiency helped to improve the photosynthetic performance of *hpr1-1* and *cat2-1*, but not *plgg1-1* (Fig. 5b). These results suggest that deficient GLYR1 activity not only improves *hpr1* but may also support mutants of the peroxisomal photorespiratory machinery more broadly.

### GLYR1 localizes to the cytosol

Since the location of GLYR1 within the cell underlies the potential mechanism by which defective GLYR1 suppresses *hpr1* and *cat2* phenotypes, we sought to definitively determine GLYR1’s subcellular localization. While several studies purport GLYR1 localization in the cytosol, these findings did not sufficiently account for the putative peroxisomal targeting signal type 1 (PTS1) tripeptide (SRE>) at GLYR1’s extreme C-terminus, which implies its potential localization in the peroxisome matrix. Apple (*Malus domestica*) GLYR1 localizes to the cytosol, but *Md*GLYR1 lacks the C-terminal putative PTS1 and therefore cannot be used to precisely infer the localization of *At*GLYR1^31^. An *At*GLYR1-GFP fusion protein transiently expressed in tobacco localized to the cytosol^30^, but again peroxisomal targeting cannot be discounted as the C-terminal PTS1 was blocked in this construct. Indirect evidence of cytosolic localization was shown in pea where the majority of glyoxylate-reducing activity was found in the cytosolic fraction, yet a small amount was detected in the chloroplast^32^. Moreover, a study using an N-terminal fluorescent tag showed the cytosolic localization of *At*GLYR1 in tobacco BY-2 and Arabidopsis cells^33^, dispelling its peroxisomal localization. However, the fluorescence signals also appeared in chloroplast-like structures in Arabidopsis mesophyll cells ^33^. Because these data did not include chlorophyll fluorescence as a marker, we cannot definitively rule out chloroplast localization of GLYR1. Therefore, it was important to obtain evidence in which *At*GLYR1’s cytosolic localization is unequivocally shown.

To this end, we generated a *35S::eYFP-GLYR1* construct and co-expressed it with the peroxisome marker mScarlet-SRL^34^ in tobacco leaves. GLYR1 appeared diffused throughout the cytosol, and no GLYR1 signal was observed to overlap with peroxisomal or chloroplast (visualized by chlorophyll autofluorescence) signals (Fig. 5c), confirming that GLYR1 is solely cytosolic.

### Transitional metabolic profiling uncovers a close link between hydroxypyruvate and GLYR1

Based on GLYR1’s mutant phenotypes, its protein localization, and the previously reported enzymatic activity, we hypothesized a photorespiratory glyoxylate shunt in the cytosol. This shunt drives the conversion of glyoxylate to hydroxypyruvate and triggers carbon flux back to the Calvin-Bensen cycle, when the primary photorespiratory pathway is deficient and when plants are exposed to high light conditions. Serine was found to accumulate in the *hpr1-1* and *cat2-1* mutants^16,35^, likely increasing the level of free serine in the cytosol. Likewise, we posited that impaired photorespiration in *hpr1-1* and *cat2-1* also causes increased glyoxylate leakage to the cytosol. The lack of a functional GLYR1 protein leads to an increase in the cytosolic level of glyoxylate, but in plants containing a functional primary photorespiratory pathway, this increase may not reach a level high enough to drive an aminotransferase activity with serine. Only in photorespiratory mutants such as *hpr1* and *cat2*, which already accumulate cytosolic glyoxylate, can GLYR1 deficiency further increase the level of cytosolic glyoxylate to a threshold to react with serine, producing hydroxypyruvate and glycine. The hydroxypyruvate generated from this reaction can then be directly catalyzed by a cytosolic HPR (such as HPR2) to produce glycerate, which returns to the Calvin-Benson cycle in the chloroplast following phosphorylation by glycerate kinase (Fig. 1). Under this hypothesis, the lack of GLYR1 function can activate a cytosolic pathway in *hpr1-1* and *cat2-1* to recycle photorespiratory carbon more efficiently back to photosynthesis, decreasing the inhibitory effects of photorespiratory intermediates and consequently improving the performance of these two mutants to compensate for the lack of HPR1 or CAT2. By contrast, loss of PLGG1 blocks the transport of glycerate into chloroplasts, preventing carbon recycling by HPR2 and therefore failing to rescue *plgg1-1*.

To test this hypothesis, we first determined how fast photorespiratory metabolites respond to high light. The partial rescue of photorespiratory metabolites observed earlier in this study (Fig. 3) was shown in plants after two weeks of growth under high light. Here we instead employed short-term treatment of photorespiratory conditions to avoid secondary effects from early metabolic events, using conditions similar to those in the RNA-seq experiment. Plants were grown under high CO_2_ and normal light conditions for 3 weeks before transfer to ambient CO_2_ and high light conditions. Plant tissue was sampled for metabolite measurements after 10 h illumination, as photorespiratory intermediates were reported to accumulate to high levels at the end of the day^23,36^.

When growing under high CO_2_, *hpr1-1* only had increased serine and slightly decreased glycerate levels (Supplementary Fig. 5), consistent with previously reported measurements of the photorespiratory intermediates under 1% (10,000 PPM) CO_2_ ^16^. The *cat2-1* mutant performed similarly to Col-0 and importantly, *glyr1-1* did not alter the levels of the photorespiratory metabolites in *hpr1-1* or *cat2-1* (Supplementary Fig. 5), indicating that the influence of *glyr1-1* is well suppressed under high CO_2_. After 10 h of ambient CO_2_ and high light conditions, the levels of 5 out of the 6 photorespiratory intermediates, including glycolate, glyoxylate, serine, hydroxypyruvate and glycerate, were significantly higher in *hpr1-1* compared to Col-0 (Fig. 6). In *cat2-1*, glycolate, serine and glycerate were significantly increased, and hydroxypyruvate also had a small but significant accumulation (Fig. 6, Supplementary Fig. 6). Interestingly, the metabolites best rescued by *glyr1-1* in both *hpr1-1* and *cat2-1* were glycolate and hydroxypyruvate (Fig. 6. Supplementary Fig. 6). Considering the enzymatic activity of GLYR1 in converting glyoxylate to glycolate, it is expected that *glyr1-1 hpr1-1* and *glyr1-1 cat2-1* have lower levels of glycolate. However, the dramatic rescue of hydroxypyruvate levels in *hpr1-1* and *cat2-1* by *glyr1-1* after only a 10-h treatment suggested that hydroxypyruvate is closely related to the function of GLYR1, which supports the cytosolic shunt we proposed. As the growth and photosynthetic phenotypes of *cat2-1* were weaker than *hpr1-1*, hydroxypyruvate accumulation in *cat2-1* was much lower and rescued better than *hpr1-1* (Fig. 6), with the hydroxypyruvate level in *glyr1-1 cat2-1* returning to wild type levels and is 22% of that in *cat2* (Supplementary Fig. 6). Glycerate abundance in the mutants showed a similar trend as that of hydroxypyruvate, although the difference between *glyr1-1 hpr1-1* and *hpr1-1* was smaller (Fig. 6), which is probably because the response of glycerate requires hydroxypyruvate as the primary responder and is therefore indirect.

**Fig. 6.**
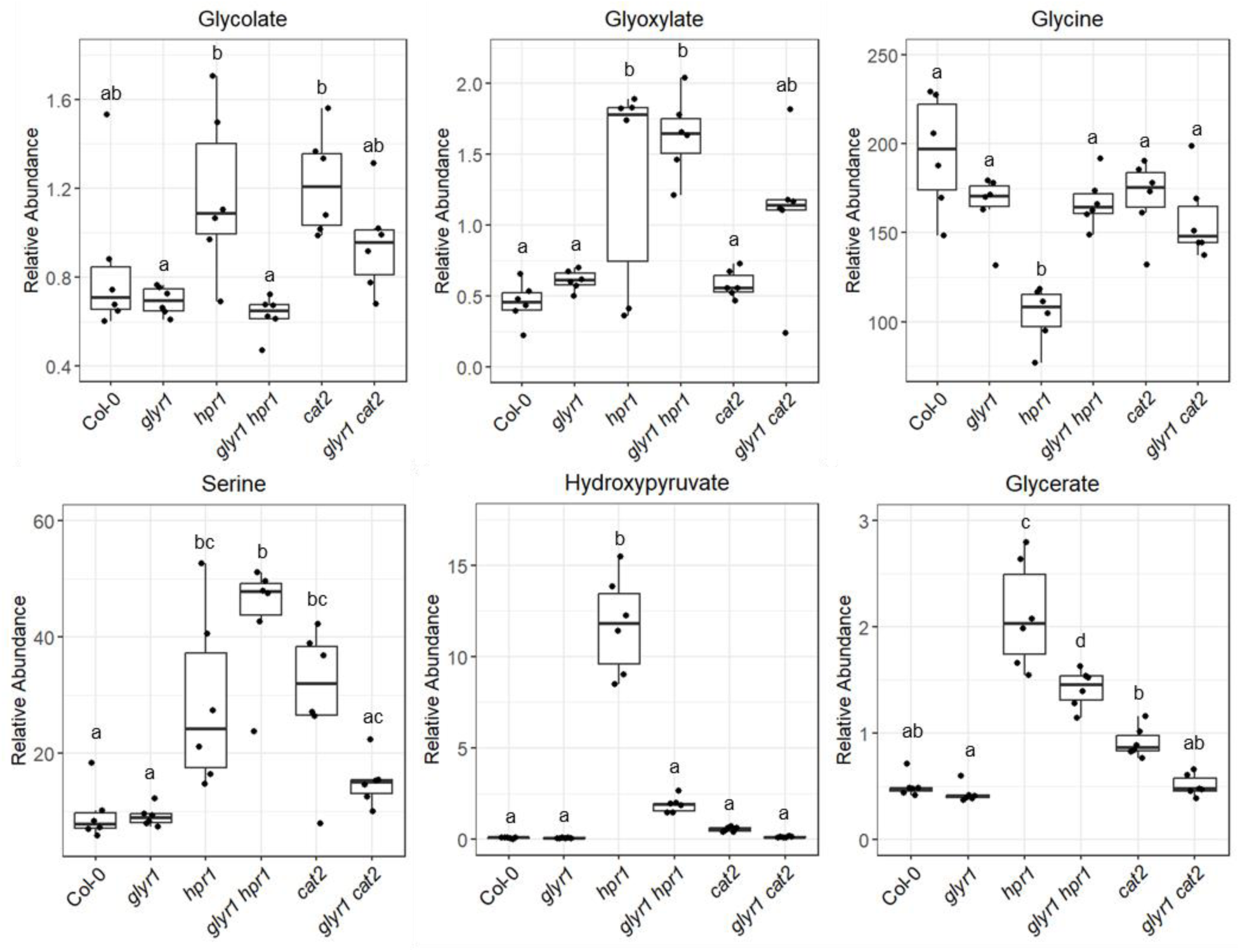
Profiling of transitional photorespiratory metabolites in plants transferred to photorespiratory environment. Plants were grown under 3 weeks of high CO_2_ and normal light, and then transferred to ambient CO_2_ and high light before lights were turned on. Leaf tissue was sampled after ∼10 h. A quick rescue of hydroxypyruvate in *hpr1* by the *glyr1* mutation was revealed. Different letters indicate statistically significant differences (p<0.05), which were determined by One-way ANOVA with Tukey’s HSD test. Biological replicates: n=6.

In summary, profiling of transitional photorespiratory metabolites suggests a close connection between hydroxypyruvate and GLYR1 under high light conditions. It supports the role of the proposed cytosolic photorespiratory shunt, in which the accumulated glyoxylate caused by defective GLYR1 in the *hpr1* and *cat2* background is converted to hydroxypyruvate.

### Feeding glyoxylate to the plant strongly inhibits growth of the wild type but can benefit *hpr1* growth

Based on our hypothesis, the availability of glyoxylate in the cytosol plays an important role in activating the non-canonical pathway in *hpr1-1*. To obtain further support for this role, we increased the level of free glyoxylate in the cytosol by directly supplying glyoxylate to the growth medium for *hpr1-1*.

Although the mechanism is still unclear, glyoxylate was reported to inhibit RuBP regeneration and Rubisco activation^37–41^ and therefore is toxic to plants. In agreement with this, wild type Col-0 plants grown in 0.4 mM glyoxylate exhibited strong growth inhibition and decreased fresh weight under normal or high light conditions (Fig. 7). However, except some suppression in root elongation (Fig. 7a), *hpr1-1* maintained similar fresh weight after glyoxylate feeding (Fig. 7b), indicating that glyoxylate can be metabolized more quickly in *hpr1-1*, possibly through the cytosolic pathway we postulated. The *glyr1-1 hpr1-1* double mutant showed a small growth inhibition upon glyoxylate treatment (Fig. 7), possibly because the total glyoxylate amount from both internal and external sources exceeds the capacity of this cytosolic shunt. In summary, results from the glyoxylate feeding experiment showed that increasing glyoxylate availability in the cytosol can benefit *hpr1-1*, which supports our hypothesis.

**Fig. 7.**
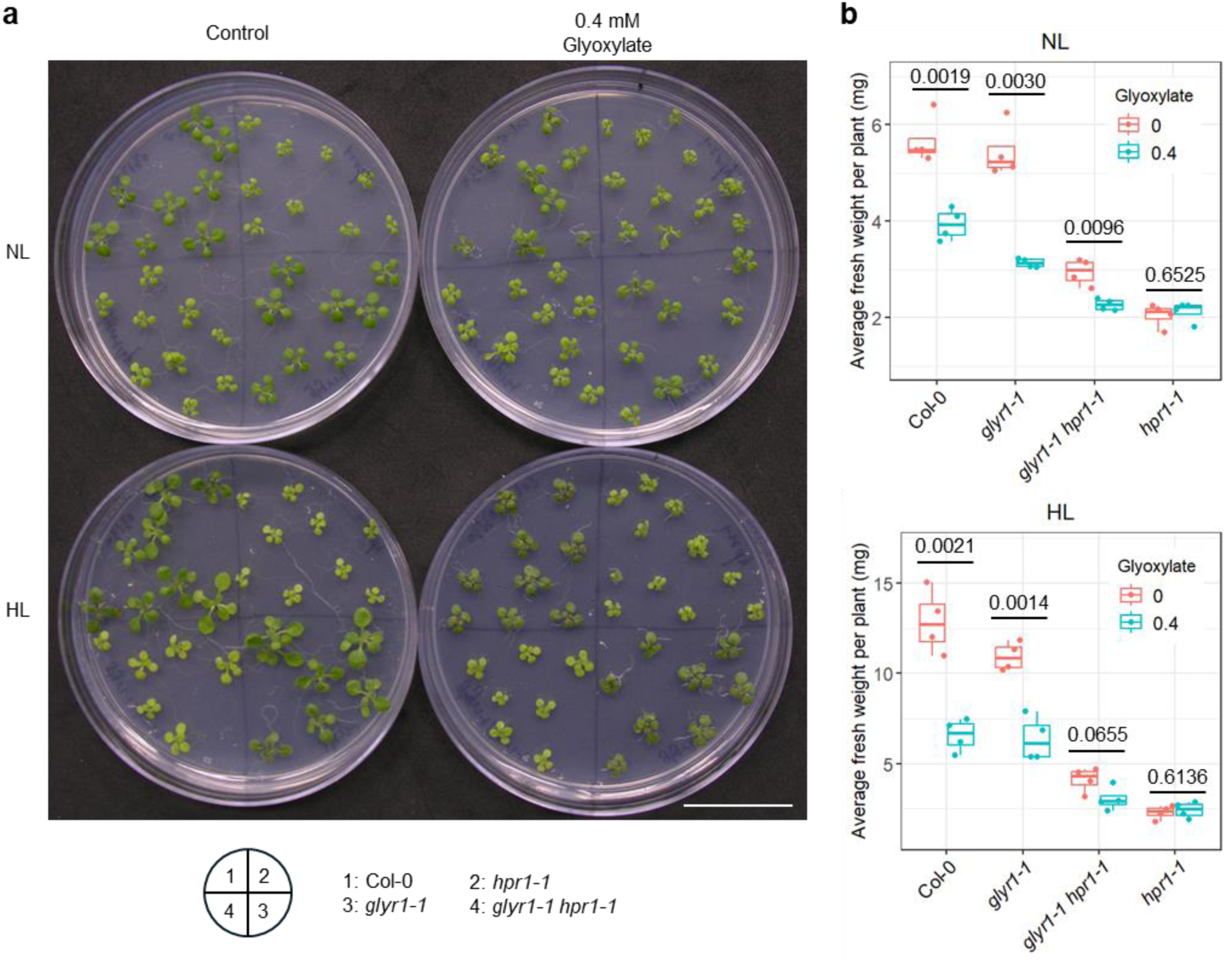
Plant growth after glyoxylate feeding. **(a)** Growth of 12-day-old seedlings on plates with or without glyoxylate under normal light (NL) or high light (HL). **(b)** Fresh weight of seedlings shown in **(a).** The *p* values determined by Student’s unpaired two-tailed *t-*test are labeled on the plot. Seedlings grown on the same plate were treated as a biological replicate. Biological replicates: n=4. Scale bars = 3 cm.

### The rescuing effect of *glyr1* in *hpr1* largely depends on HPR2

Given that our proposed photorespiratory glyoxylate shunt requires the activity of cytosolic HPR, we investigated the role of HPR2 in this shunt because it has a stronger role in photorespiration than HPR3^17^. Null mutant *hpr2-3* (Supplementary Fig. 7) was used to generate a triple knockout line *glyr1-1 hpr1-1 hpr2-3*. Because mutants defective in both *HPR1* and *HPR2* genes are already stunted, grow poorly in ambient air, and are intolerant to high light treatment, we first grew all the lines under high CO_2_ and normal light conditions, during which all mutants showed comparable morphologies as Col-0 (Fig. 8a, Day 0). After three weeks and at the end of the dark period, plants were moved to ambient CO_2_ and high light conditions, where they were kept for 9 days. This treatment led to much smaller rosettes in *hpr1-1* compared to Col-0, which was partially rescued in *glyr1-1 hpr1-1* and the original suppressor *shpr7* (Fig. 8a). At Day 9, although *hpr1-1 hpr2-3* and *glyr1-1 hpr1-1 hpr2-3* performed poorly compared with other lines, they looked similar with each other, supporting our conclusion that, at least at this time point, *glyr1-1*’s role in helping carbon recycle back to the chloroplast largely depends on a functional HPR2. At Day 18, *glyr1-1 hpr1-1 hpr2-3* became slightly bigger than *hpr1-1 hpr2-3* (Fig. 8a), indicating that *glyr1-1* may also act through other proteins (such as HPR3) for its role under longer photorespiratory conditions.

**Fig. 8.**
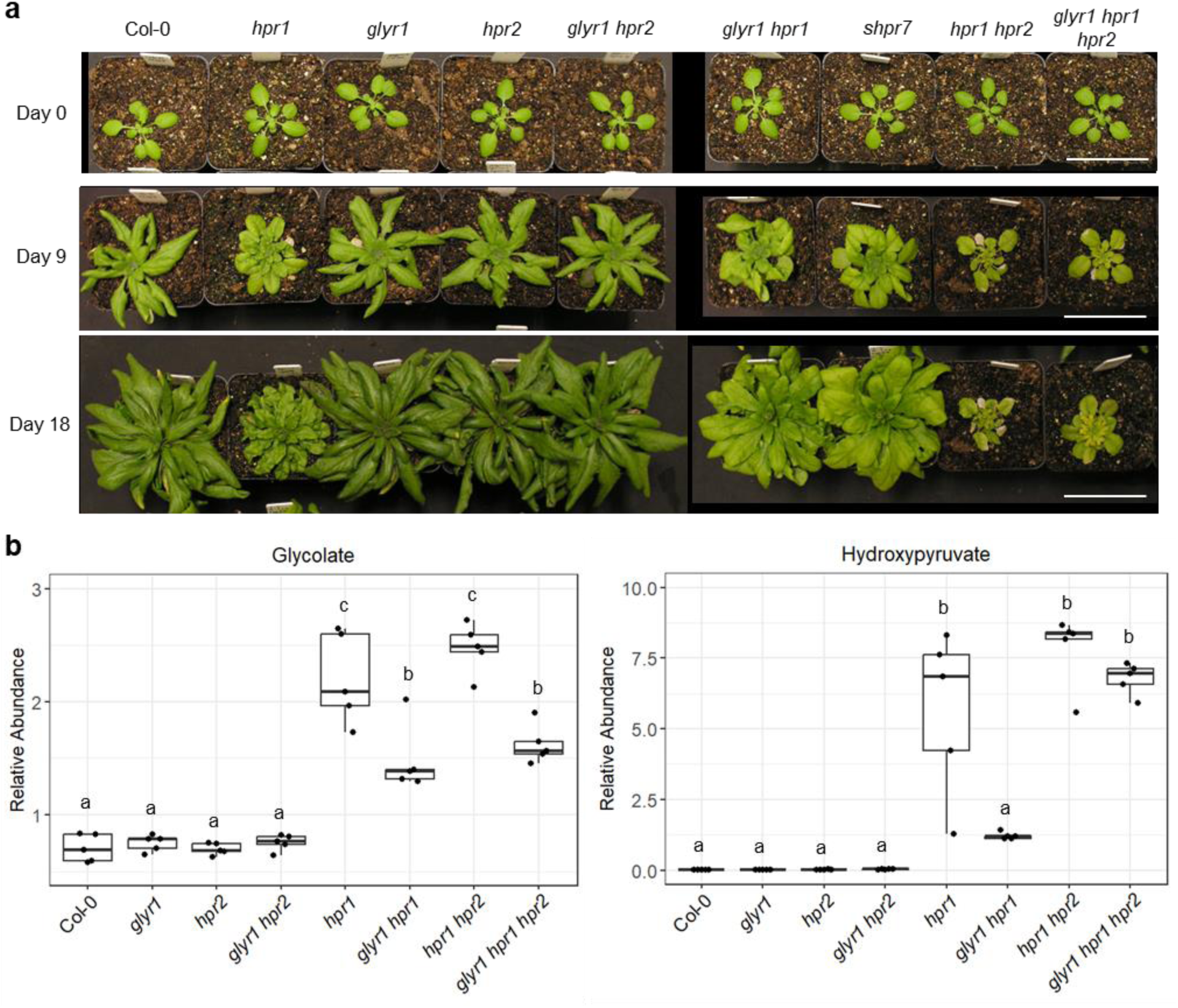
The role of defective GLYR1 in suppressing *hpr1* is largely dependent on HPR2. **(a)** Plants grown under 3 weeks of high CO_2_ and normal light before being transferred to ambient CO_2_ and high light on Day 0. Scale bars = 5 cm. **(b)** Glycolate and hydroxypyruvate levels at the transitional stage. Plants were grown under 3 weeks of high CO_2_ and normal light, and then transferred to ambient CO_2_ and high light before the lights were on. Leaf tissue was sampled after ∼10 h. Different letters indicate statistically significant differences (p<0.05), which were determined by One-way ANOVA with Tukey’s HSD test. Biological replicates: n=5.

Profiling of transitional photorespiratory metabolites during the transfer to photorespiratory conditions was also performed on mutants grown under high CO_2_ and normal light for 3 weeks followed by 10-h treatment with ambient CO_2_ and high light. Here we focused on glycolate and hydroxypyruvate, the two photorespiratory metabolites exhibiting fast changes in *glyr1 hpr1* and *glyr cat2* in comparison to *hpr1* and *cat2*, respectively (Fig. 6, Supplemental Fig. 6). Similar to the rescued glycolate level in *glyr1-1 hpr1-1*, *glyr1-1 hpr1-1 hpr2-3* also showed a lower glycolate level than *hpr1-1 hpr2-3* (Fig. 8b), confirming that the glycolate production via GLYR1 is greatly lost. By contrast, the level of hydroxypyruvate was not significantly changed in *glyr1-1 hpr1-1 hpr2-3* (Fig. 8b), supporting the importance of HPR2 in this cytosolic pathway.

## Discussion

In this study, we provided evidence for a cytosolic glyoxylate shunt of photorespiration that can improve carbon recycling in the absence of a functional major photorespiratory pathway under high light, supporting the metabolic flexibility of the photorespiration network. Under high light, deficiency in GLYR1, a cytosolic enzyme that converts glyoxylate to glycolate, can partially rescue the phenotypes of the photorespiration mutants *hpr1* and *cat2* but not those of the glycolate/glycerate transporter mutant *plgg1*. Further investigations showed that hydroxypyruvate is closely connected to GLYR1’s function, and the availability of glyoxylate and HPR2 in the cytosol is important for the suppression. These results led us to propose a novel cytosolic photorespiratory shunt, which seems to be critical when the major photorespiratory pathway is compromised and when plants are exposed to conditions under which high photorespiratory fluxes are needed, such as high light. Specifically, a reaction between glyoxylate and serine is catalyzed by an aminotransferase to produce hydroxypyruvate and glycine (Fig. 1). Via HPR2, cytosolic hydroxypyruvate is subsequently converted to glycerate that eventually returns to the Calvin-Benson cycle, thus reducing the accumulation of the toxic photorespiratory intermediates and enhancing carbon recycling (Fig. 1). We predict that, in wild type plants, this shunt may only be activated under extremely high photorespiratory fluxes.

Plants contain two GLYR proteins, the cytosolic GLYR1 and the plastid/mitochondrion-dual localized GLYR2, both of which are believed to be involved in aldehyde detoxification during abiotic stress ^29,42,43^. Although GLYR1 was shown to be more efficient in its glyoxylate reductase activity, it can also convert succinic semialdehyde to γ-hydroxybutyrate in γ-aminobutyrate (GABA) metabolism^28,29^. During abiotic stress, the expression of both *GLYR1* and *GLYR2* is upregulated, presumably to help detoxify the accumulated succinic semialdehyde in plants^29,42,43^. However, the positive role of succinic semialdehyde reduction during abiotic stress contradicts the rescuing effect of *glyr1* in *hpr1* under high light, and GABA metabolism is not known to be directly linked to photorespiration. Therefore, the succinic semialdehyde reductase activity of GLYR1 does not seem to be involved in its role related to HPR1 and CAT2.

Since the knockout mutants of *GLYR1* are comparable to Col-0 in plant growth and levels of the photorespiratory metabolites, this cytosolic photorespiratory shunt may not be important when the main photorespiratory pathway in the peroxisome is functional, possibly due to the predominant peroxisomal location of glyoxylate. Glyoxylate is highly reactive and can inhibit photosynthesis and other metabolic reactions, thus is toxic to the plant^37–41^. There are two major glyoxylate metabolic pathways in the peroxisome to maintain its homeostasis: the main photorespiratory pathway in photosynthetic tissue and the glyoxylate cycle in seeds in which glyoxylate is produced and catabolized ^44,45^. Further, the cytosolic GLYR1 and the plastidic/mitochondrial GLYR2 were believed to scavenge the glyoxylate leaked from the peroxisome^30,44^. Since cytosolic glyoxylate may become toxic to plants, it is possible that the cytosolic shunt is activated only when the main peroxisomal pathway is defective and under conditions that need very high photorespiratory flux.

We provided evidence that HPR2 is crucial to converting cytosolic hydroxypyruvate to glycerate in the photorespiratory glyoxylate shunt, as the *glyr1 hpr1 hpr2* triple mutant maintains 86% of the hydroxypyruvate content observed in *hpr1 hpr2* (Fig. 8b). However, the minor decrease in hydroxypyruvate content, as well as the slightly bigger *glyr1 hpr1 hpr2* rosette size compared to *hpr1 hpr2* after 18 days of high light treatment (Fig. 8a) suggests the existence of other mechanisms in cytosolic hydroxypyruvate reduction independent of HPR2. Although the role of HPR3 in reducing hydroxypyruvate is minor, knocking out *HPR3* further exacerbates the photorespiratory phenotypes of the *hpr1 hpr2* double mutant^17^. Therefore, HPR3 may be involved in maintaining this photorespiratory shunt in the cytosol when both HPR1 and HPR2 are absent.

We also hypothesize that there is a cytosolic aminotransferase with similar activity as the peroxisomal serine:glyoxylate aminotransferase (SGAT). Although SGAT activity was only detected in the peroxisome in previous studies^46,47^, cytosolic activity may not be detectable unless the cytosolic route is activated. SGAT does not have any apparent homologs in the Arabidopsis genome^47^, but substrate promiscuity is common for aminotransferases^48^. SGAT is also known as AGT1 because of its alanine:glyoxylate transferase activity; additionally, it can catalyze other amino donor:acceptor combinations such as serine:pyruvate and asparagine:glyoxylate^46,48^. Human (*Homo sapiens*) AGT2, which localizes to mitochondria and has broad substrate specificity^48^, has three homologs in Arabidopsis: *At*AGT2, *At*AGT3, and *At*PYD4 (pyrimidine 4)^48,49^. *At*AGT2 was reported to localize to mitochondria and peroxisomes^50^. The enzymatic activity and localization of *At*AGT3 and *At*PYD4 are unclear, thus these proteins may be candidates for the hypothetical cytosolic aminotransferase with SGAT activity.

Some of the metabolic profiles of glycine and serine in this work have complicated patterns, possibly due to their participation in other metabolic pathways. Under ambient (21%) O_2_ conditions, 32% of the photorespiratory carbon is thought to leave this pathway as serine^51,52^, indicating that photorespiratory serine may be strongly connected with other metabolisms. Serine has also been shown to play a role in the biosynthesis of other amino acids, proteins, and lipids^53^. While glycine is converted to serine in the mitochondrion during photorespiration, serine can also be converted to glycine in the cytosol and plastid^54^. This serine-glycine interconversion is an important component in one-carbon metabolism, which is crucial for synthesizing nucleotides and methylated compounds and maintaining nutrient balance^54,55^. Furthermore, it has been reported that excess glycine from photorespiration is used for protein synthesis^56^ and glutathione accumulation^57^. Finally, glycine and serine were found in the vacuole^58,59^, which may represent an inactive pool or have a slow response to environmental changes.

Our work has provided evidence for the important role of an alternative photorespiratory route in the cytosol under high light conditions when the major photorespiratory pathway is defective, supporting the metabolic flexibility of photorespiration and substantiating the contribution of photorespiration to stress response. Further investigations are needed to obtain a more comprehensive understanding of the regulation of photorespiration, which may ultimately help to generate new crop varieties with high productivity without compromising their stress tolerance.

## Methods

### Plant materials and growth conditions

Wild-type and mutant lines of *Arabidopsis thaliana* used in the study are all from the ecotype Col-0. T-DNA insertion mutants were obtained from the Arabidopsis Biological Resource Center (ABRC, Ohio State University, USA) and confirmed by PCR-based genotyping. Previously characterized mutants are *hpr1-1* (SALK_067724)^16^, *cat2-1* (SALK_076998)^60^, *plgg1-1* (SALK_053469)^36^, and *glyr1-1* (SALK_057410)^29^. Previously uncharacterized mutants used in this study are *glyr1-2* (SALK_202680C), *glyr1-3* (SALK_203580C), and *hpr2-3* (SALK_105876). Primers used for genotyping are listed in Supplementary Table 1.

Arabidopsis seeds were sown on plates containing half-strength Linsmaier and Skoog basal salt (1/2 LS, Caisson Labs), 1% sucrose and 0.8% agar (Phytoblend, Caisson Labs). After seed stratification in the dark at 4 °C for 3 to 7 days, plates were placed in the Percival Intellus Environmental Controller under normal light (∼100 μmol m^-2^ s^-1^ white light), 21 °C, and 12h/12h light/dark cycle. At ∼1.5 weeks old, seedlings were transplanted to the soil and moved to growth chambers with the same growth conditions. For high light treatment, 2-week-old plants were transferred to a chamber with ∼700 μmol m^-2^ s^-1^ white light, and the same temperature and photoperiod as those for growth in normal light. Plants grown under high CO_2_ were directly grown in the soil under 0.2% CO_2_, with the same settings for other parameters.

For the glyoxylate feeding experiment, glyoxylate was filter-sterilized and added to the autoclaved medium to a final concentration of 0.4 mM. Plates were placed in Percival with normal light and growth chamber with high light, respectively, at the same time.

### Suppressor screening and mapping

Seeds of *hpr1-1* were treated with ethyl methanesulfonate (EMS) to induce random mutations. M_1_ plants were self-pollinated and M_2_ seeds were harvested. M_2_ plants were grown under high light for suppressor screening, and individuals that grew bigger or greener than *hpr1-1* were selected. Candidates with persistent suppressor phenotypes in the M_3_ generation were backcrossed to *hpr1-1*. Plants displaying the suppression phenotypes in the BC_1_F_2_ generation were used for additional backcrosses.

In the BC_3_F_2_ generation of the *shpr7* suppressor, 80 individuals with the suppression phenotype were selected. Genomic DNA from these plants was extracted by Wizard Genomic DNA Purification Kit (Promega) and pooled together as one sample, which was used for DNBSEQ PE150 whole genome sequencing with 80X coverage. To minimize the influence of results by unrelated background mutations, DNA was extracted from 29 *hpr1-1* individuals and pooled together for whole genome sequencing with 30X coverage. Sequencing and bioinformatic analysis were performed by BGI Genomics (https://www.bgi.com). Selfed progenies from BC_3_F_2_ plants exhibiting suppression phenotypes were used for follow-up experiments.

### Generation of transgenic lines and RT-PCR

The *35S::FLAG-GLYR1* construct was generated by Gateway cloning according to manufacturer’s instructions (ThermoFisher). Briefly, the coding sequence of *GLYR1* was amplified from cDNA from Col-0 and cloned into the entry vector pDONR207 through the BP reaction. Next, GLYR1 was inserted into the destination vector pEarleyGate202 through the LR reaction. Entry and expression constructs were confirmed by Sanger sequencing. The construct was transformed into *Agrobacterium tumefaciens* strain GV3101 by electroporation, and Arabidopsis lines were transformed by *Agrobacteria* using floral dipping. Positive transformants were selected on 1/2 LS plates containing 1% sucrose and 10 μg/ml glufosinate ammonium.

For RT-PCR analysis, RNA was extracted from the leaf tissue using NucleoSpin RNA Plant kit (MACHEREY-NAGEL), then reverse transcribed using the High-Capacity cDNA Reverse Transcription Kit (Applied Biosystems). The coding sequence of the target genes were PCR-amplified from cDNA using the GoTaq Green Master Mix (Promega) and gene-specific primers (Supplementary Table 1).

### Protein preparation and immunoblot analysis

Whole rosettes from 4-week-old plants, which had been grown for 2 weeks under normal light followed by 2-week growth under high light, were collected and ground in liquid nitrogen, after which ∼60 mg of the powder was homogenized with 500 μl Extraction Buffer [150 mM Tris-HCl (pH 6.8), 7.5% β-mercaptoethanol, 3% sodium dodecyl sulfate, and half a cOmplete mini protease inhibitor tablet (Roche)]. Samples were centrifuged at 17,000xg at 4 °C for 10 min, and the supernatants were transferred to new tubes.

A previously described method was used for immunoblotting^61^. Specifically, protein samples were combined with 4X Laemmli Buffer and boiled at 95 °C for 5 min. 40 μl samples were loaded onto a 4-20% Mini-PROTEAN TGX pre-cast gel (Bio-Rad) with the Dual Color Protein Ladder (Bio-Rad). After separation on the gel, proteins were transferred to a nitrocellulose membrane using Power Blotter XL System (Invitrogen) at 25V for 7 min, which was then blocked with 5% milk in TBST for 1 h. The anti-GLYR1 antibody was developed by PhytoAB Inc. (CA, USA), based on the antigen peptide PAFPLKHQQKDMRLA with an additional C at the C-terminus for conjugation. The membranes were incubated in 1:1000 rabbit anti-GLYR1 antiserum in the blocking buffer overnight at 4 °C, washed multiple times with TBST, incubated in 1:10,000 goat anti-rabbit IgG HRP secondary antibody (PhytoAB Inc.) in TBST for 2 h, washed again with TBST, and then developed with the SuperSignal West Pico PLUS Chemiluminescent Substrate kit (Thermo Scientific). A replicated protein gel with the same loading and running conditions was used for Coomassie staining.

### Metabolite extraction and quantification

Leaf samples were collected in the late afternoon, after ∼10 h light treatment. Two to three well-expanded leaves with similar age were taken from plants with similar morphologies. In the cases where the mutants show different morphologies, the whole rosettes were taken. Plant tissue was frozen and ground in liquid nitrogen.

Metabolite extraction was performed using the methanol/chloroform/water method and quantification was conducted by Gas Chromatography–Mass Spectrometry (GC-MS), as described previously^51^. Specifically, 500 μl chloroform/methanol (3:7, v/v) was added to the tissue powder, followed by incubation at -20 °C for 2 h with occasional shaking. Then the internal standard adonitol and 400 μl water were added to the sample, followed by vigorously mixing and centrifugation at 4 °C, 12000 xg for 10 min. The upper phase (methanol in water) was taken, dried in a lyophilizer and stored in -80°C.

Before analysis, samples were derivatized by adding 20 mg/ml methoxyamine hydrochloride dissolved in dry pyridine and incubated at room temperature overnight. Next, the reaction mixture was silylated by N, O-Bis (trimethylsilyl) trifluoroacetamide with 1% trimethylchlorosilane at 60°C overnight. The trimethylsilyl (TMS) derivatives were analyzed by an Agilent 7890A GC system/5975C inert XL Mass Selective Detector with a 30 m VF-5ms column (Agilent). 1 μl of each sample was injected into the 230°C inlet, where splitless, 10:1 split, or 20:1 split mode was chosen depending on the levels of the metabolites in plant samples. The oven temperature gradient is as follows: 40 °C for 1 min, increased to 80 °C in 1 min, then further increased by 10 °C/min to 240 °C followed by 20 °C /min to 320 °C, and 5 min holding at 320 °C. Scan mode of the MS was used to monitor ions with mass to charge ratio (m/z) between 50–600.

Metabolites were identified by m/z values, using retention time in comparison with authentic standards or the NIST Mass Spectral Library. Software MassLynx (Waters) was used for peak extraction and integration. Metabolites were quantified against the internal standard.

### Photosynthetic measurement

Using a previously reported method^10^, the Dynamic Environment Photosynthesis Imager (DEPI)^62^ was used to measure the quantum efficiency of photosystem II (Φ_II_) of 2-week-old seedlings grown in soil under normal light at 21 °C and 12h/12h light/dark cycle. Φ_II_ was calculated from (Fm′-Fs)/Fm′, where Fs is the chlorophyll fluorescence emission from light-adapted leaf at steady state and Fm’ is the maximum fluorescence from light-adapted leaf during a saturating pulse of light. Custom software (Visual Phenomics) was used to process the fluorescence images^63^, and heatmaps were generated with the software OLIVER (https://caapp-msu.bitbucket.io/projects/oliver/index.html).

### Tobacco infiltration and confocal microscopy

GLYR1 was N-terminally tagged with eYFP using the Gateway LR Clonase II Enzyme mix (ThermoFisher) by combining entry vector pENTR223-GLYR1 (ABRC, G82382) and pEarleyGate104, according to manufacturer’s instructions. The expression vector (eYFP-GLYR1) was confirmed by Sanger sequencing and transformed into GV3101 *Agrobacterium tumefaciens* cells by electroporation.

5 ml Agrobacteria containing the peroxisome marker (mScarlet-SRL in pGWB2)^34^ and eYFP-GLYR1 were grown in Luria broth (LB, Rif_25_/Gent_10_/Kan_50_) with 225 rpm shaking at 28 °C overnight. Separate flasks containing 25 ml LB (Rif_25_/Gent_10_/Kan_50_) with 100 μM acetosyringone were inoculated with 1 ml of the overnight cultures and grown at 28 °C to OD_600_ between 0.6 and 0.8. The cells were harvested by centrifugation at 3000 xg for 10 min and then resuspended in MMA infiltration buffer (10 mM MES pH 5.7, 10 mM MgCl_2_, 100 μM acetosyringone) to adjust the OD_600_ to 0.6. The cell resuspensions were then incubated at room temperature (∼22 °C) for 1 h. The peroxisome marker and eYFP-GLYR1 cultures were combined in 1:1 volume and infiltrated into 6-week-old *Nicotiana tabacum* leaves with a 1 ml needleless syringe. Tobacco plants were recovered in a 12h/12h photoperiod chamber for 2 days before imaging.

Infiltrated tobacco leaves were imaged with an Olympus Fluoview 1000 spectral-based confocal laser scanning microscope with a 40X oil objective (NA: 1.30). Images were captured using the following excitation and emission parameters: eYFP-GLYR1 (ex: 515 nm, em: 530-560 nm), mScarlet-SRL (ex: 559 nm, em: 580-615 nm), and chlorophyll autofluorescence (ex: 515 nm, em: 655-755 nm).

### RNA sequencing and data analysis

After growing under high CO_2_ and normal light for 3 weeks, plants were moved to ambient CO_2_ at the end of the dark period, followed by 3 h or 10 h high light treatment. Five biological replicates of each genotype (Col-0, *hpr1*, *glyr1*, *shpr7*, *glyr1 hpr1*) were used, and 2-3 well-expanded leaves of similar age were sampled on each plant. Plant tissue was frozen and ground in liquid nitrogen.

Total RNA was isolated using the NucleoSpin RNA Plant kit (MACHEREY-NAGEL), after which DNA was further depleted using the TURBO DNA-free Kit (Invitrogen). RNA-seq was performed by the MSU Genomics Core, who prepared mRNA libraries with the KAPA mRNA HyperPrep Kit (Roche) and pooled them together with samples from other researchers onto one S4 lane of Illumina NovaSeq 6000. The sequencing yielded 2×150 bp paired end reads with ∼20M read pairs per sample.

Data analysis was performed by the MSU Bioinformatics Core. The nf-core/rnaseq v3.10.1 pipeline (https://zenodo.org/records/7505987) built with Nextflow v22.10.4^64^ was used to process and quantify transcriptomic reads using the standard defaults unless otherwise specified. Briefly, Salmon v1.9.0^65^ and the fq v0.9.1 (https://github.com/stjude-rust-labs/fq) were used to sub-sample FastQ files and auto-infer read strandedness. Adapter and quality trimming were performed using Trim Galore! v0.6.7 (https://zenodo.org/records/5127899) with Cutadapt v3.4^66^. STAR v2.7.9a was used to map the raw FastQ reads to the reference genome and project the alignments onto the transcriptome^67^. The alignments were sorted and indexed using SAMtools v1.16.1^68^, and downstream BAM-level transcript quantification was performed with Salmon v1.9.0 with the --seqBias --gcBias tags^65^. Reads were mapped to the TAIR 10.1 version of the *A. thaliana* genome (GCA_000001735). Note that genes with missing transcript IDs were filtered from the GTF file, which is a known issue with some GTF files (see https://github.com/nf-core/rnaseq/issues/1086).

DESeq2 v1.38.3 was used to perform differential gene expression analysis for 3- and 10-h samples separately^69^. Tximport v1.26.1 was used to import transcript abundances and construct a gene-level DESeqDataSet object from Salmon quant.sf files^70^. Genes were filtered for those with a count of at least 10 in 5 samples.

Heatmaps of differentially expressed genes, photosynthetic, and photorespiration genes were generated using the pheatmap v1.0.12 package (https://github.com/raivokolde/pheatmap). Genes (rows) were clustered using row scaled, Euclidean distance and the ward.D clustering approach.

### Statistics and plots

Data analysis was performed using Microsoft Excel and Rstudio. Student’s unpaired two-tailed *t*-test was used for pairwise comparison. One-way ANOVA with Tukey’s HSD test was used for multi-comparison. Boxplot center lines show the medians, and the lower and upper hinges correspond to the first and third quartiles. Whiskers extend from the hinges to the largest or smallest values that are no further than 1.5 times of the interquartile range. Data points are represented as individual dots.

## Acknowledgements

We thank Drs. Stefanie Tietz and Jiying Li for performing EMS mutagenesis and the initial screening; David Hall and Dr. David Kramer for helping with experiments using the Dynamic Environment Photosynthesis Imager and providing insightful thoughts; Cassandra Johnny (MSU Mass Spectrometry and Metabolomics Core) and Drs. Xinyu Fu and Kelem Alamrie for technical support in metabolic measurements; Dr. Nicholas Panchy (MSU Bioinformatics Core) for analyzing the RNA-seq data; Drs. John Froehlich and Sarah Stainbrook for technical support in immunoblotting; and Linnea Hartz, Joy Li, and Carter Pasternak for technical help. This work was supported by the Chemical Sciences, Geoscience and Biosciences Division, Office of Basic Energy Sciences, Office of Science, U.S. Department of Energy (DE-FG02-91ER20021) and The National Science Foundation (MCB 2148206) to J.H.

## Author contributions

J.H. conceived the study. X.J. and J.H. designed the experiments with input from B.J.W. and A.M.K. X.J. performed most of the experiments with technical support from B.J.W. and A.M.K. A.M.K. performed GLYR1 localization and RNA-seq data analysis. X.J., A.M.K. and J.H. cowrote the paper with valuable contributions from B.J.W.

## Competing interests

The authors declare no competing interests.

**Correspondence** and requests for materials should be addressed to Jianping Hu.

## Notes

### Competing Interest Statement

The authors have declared no competing interest.

