## Supplemental Material for "A photorespiratory glyoxylate shunt in the cytosol supports photosynthesis and plant growth under high light conditions in Arabidopsis"

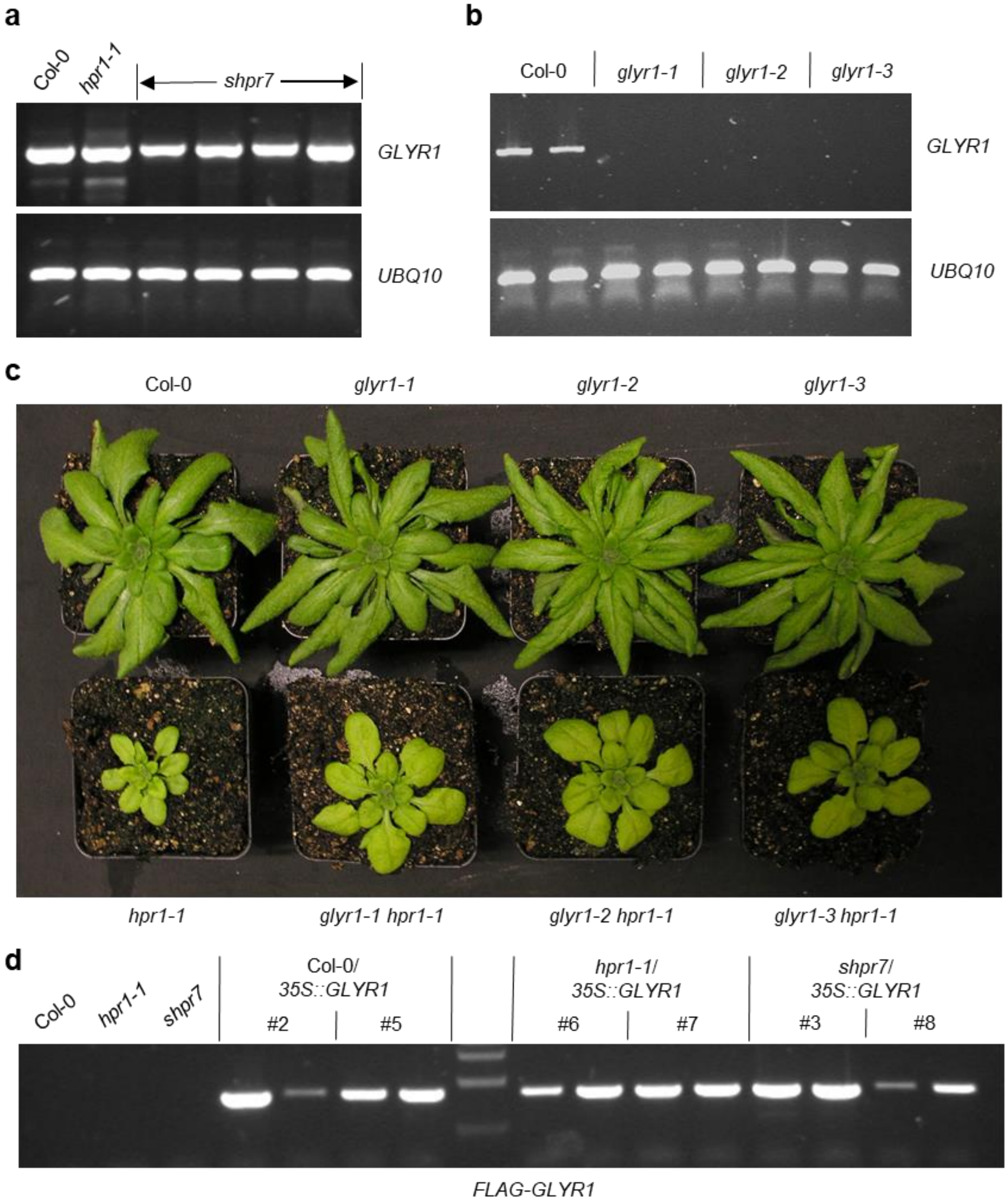


**Supplementary Fig. 1. Characterizations of *shpr7*, T-DNA insertion lines of *GLYR1*, and *GLYR1*-overexpression lines. (a)** *GLYR1* expression in *shpr7* plants grown under 2-week normal light followed by 4.5-week high light. **(b)** *GLYR1* expression in the T-DNA insertion lines grown under 2-week normal light followed by 3-week high light. *UBQ10* was used as a control in **(a)** and **(b)**. **(c)** The *glyr1* single mutants and *glyr1 hpr1* double mutants grown under 2-week normal light followed by 2.5-week high light. **(d)** *FLAG-GLYR1* expression in transgenic lines grown under 2-week normal light followed by 2.5-week high light.

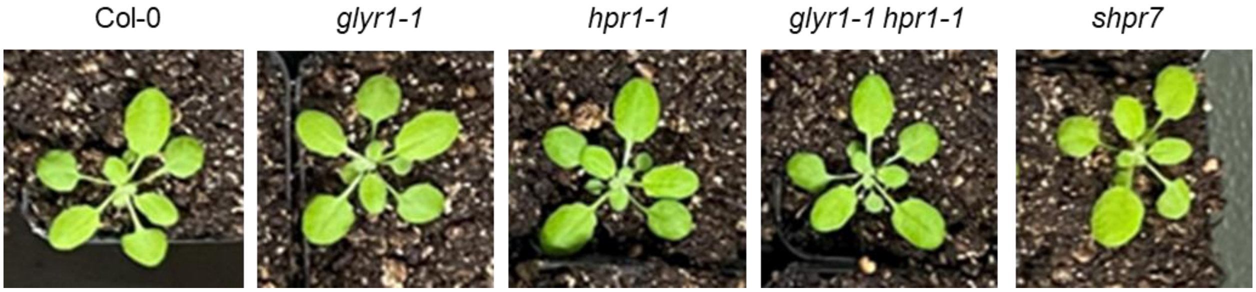


**Supplementary Fig. 2. Appearance of plants grown under high $CO_2$ conditions.** Plants were grown under high $CO_2$ and normal light for 3 weeks.

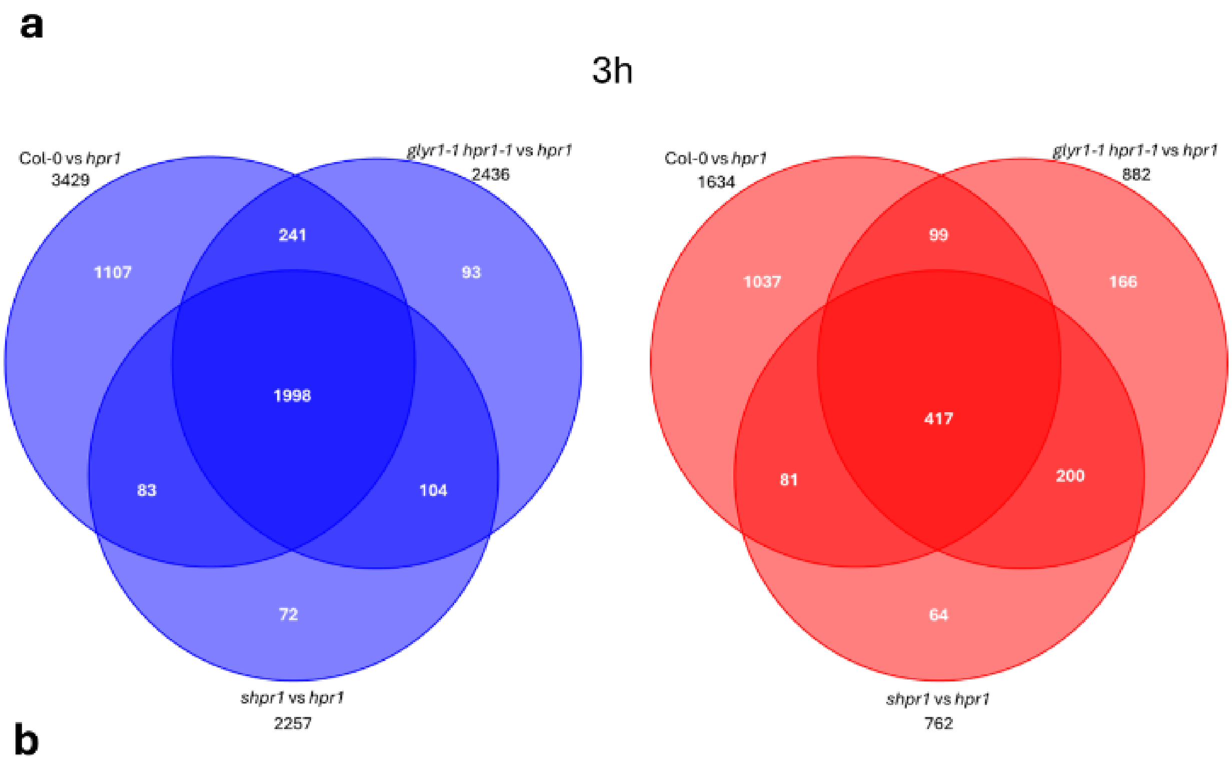


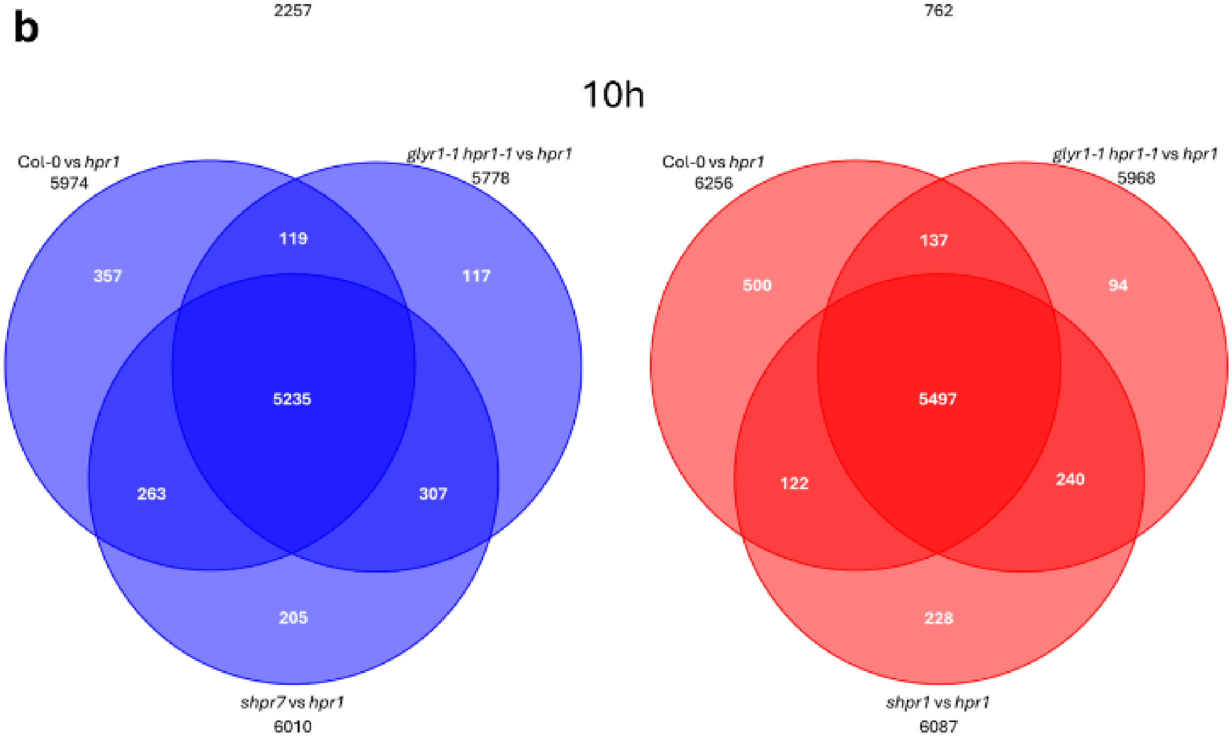


**Supplementary Fig. 3. Venn diagrams of the differentially expressed genes (DEGs).** DEGs with |Log2Fold-Change|>1 and $p_{adj}<0.01$ after 3 h **(a)** and 10 h **(b)** of plant transfer to photorespiratory conditions are shown for both downregulated (blue) and upregulated (red) genes in Col-0, *glyr1-1 hpr1-1*, and *shpr1* compared to *hpr1-1*. The majority of the DEGs in Col-0 compared to *hpr1-1* show the same regulation pattern in *glyr1-1 hpr1-1* and *shpr7* compared to *hpr1-1*, indicating that deficient GLYR1 restores the *hpr1* transcriptome to wild type expression.

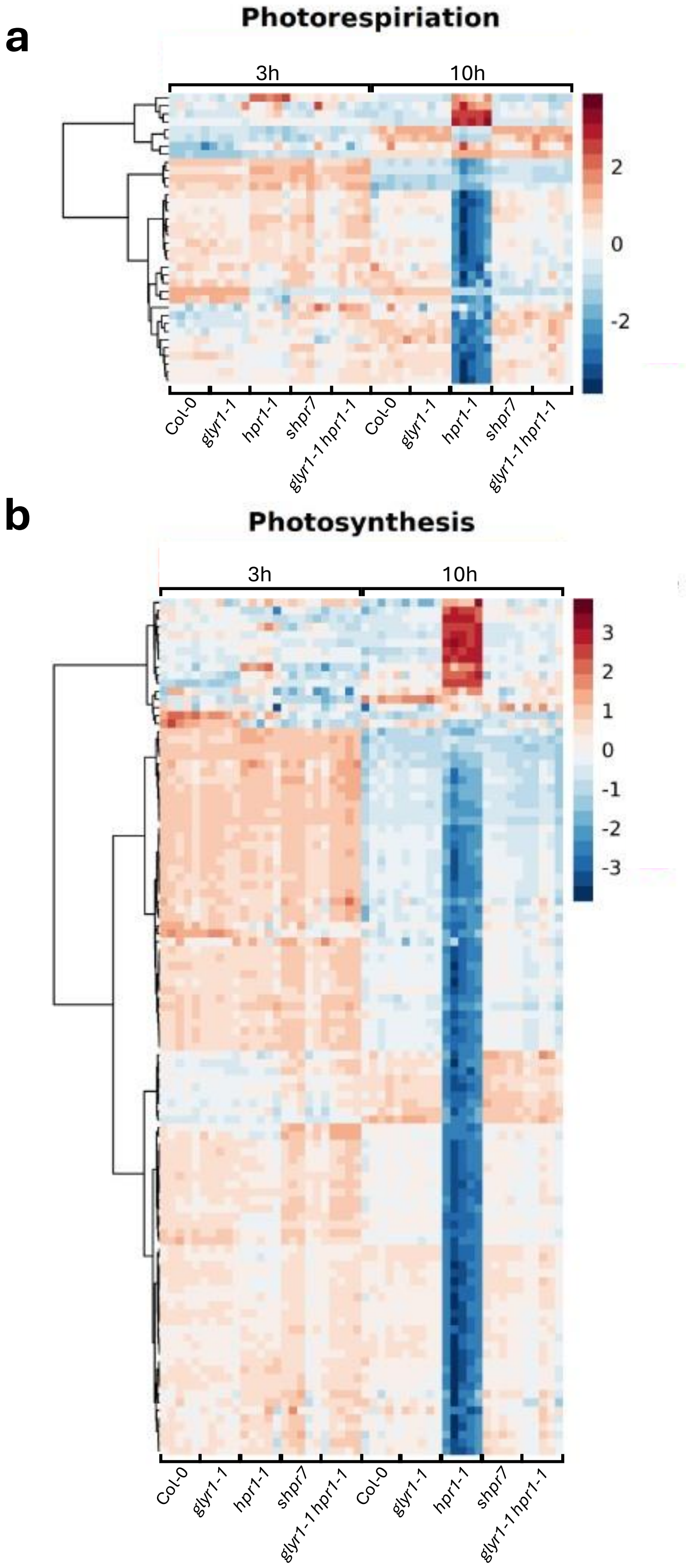
a
Photorespiriation
3h
10h
2
0
-2
Col-0
glyr1-1
hpr1-1
shpr7
glyr1-1 hpr1-1
Col-0
glyr1-1
hpr1-1
shpr7
glyr1-1 hpr1-1
b
Photosynthesis
3h
10h
3
2
1
0
-1
-2
-3
Col-0
glyr1-1
hpr1-1
shpr7
glyr1-1 hpr1-1
Col-0
glyr1-1
hpr1-1
shpr7
glyr1-1 hpr1-1

**Supplementary Fig. 4. Heatmaps of the expression profiles of genes related to photorespiration (a) and photosynthesis (b)**. Genes with ≥4-fold change (|Log2Fold-Change|>2 and $p_{adj}<0.01$) in Col-0 vs. *hpr1-1* in all genotypes at 3 h and 10 h of treatment with photorespiratory conditions are shown. The transcriptional reprogramming of photorespiration- and photosynthesis-related genes in *hpr1-1* is rescued by deficient GLYR1, as the expression patterns are reverted to Col-0 levels in *glyr1-1 hpr1-1* and *shpr7*.

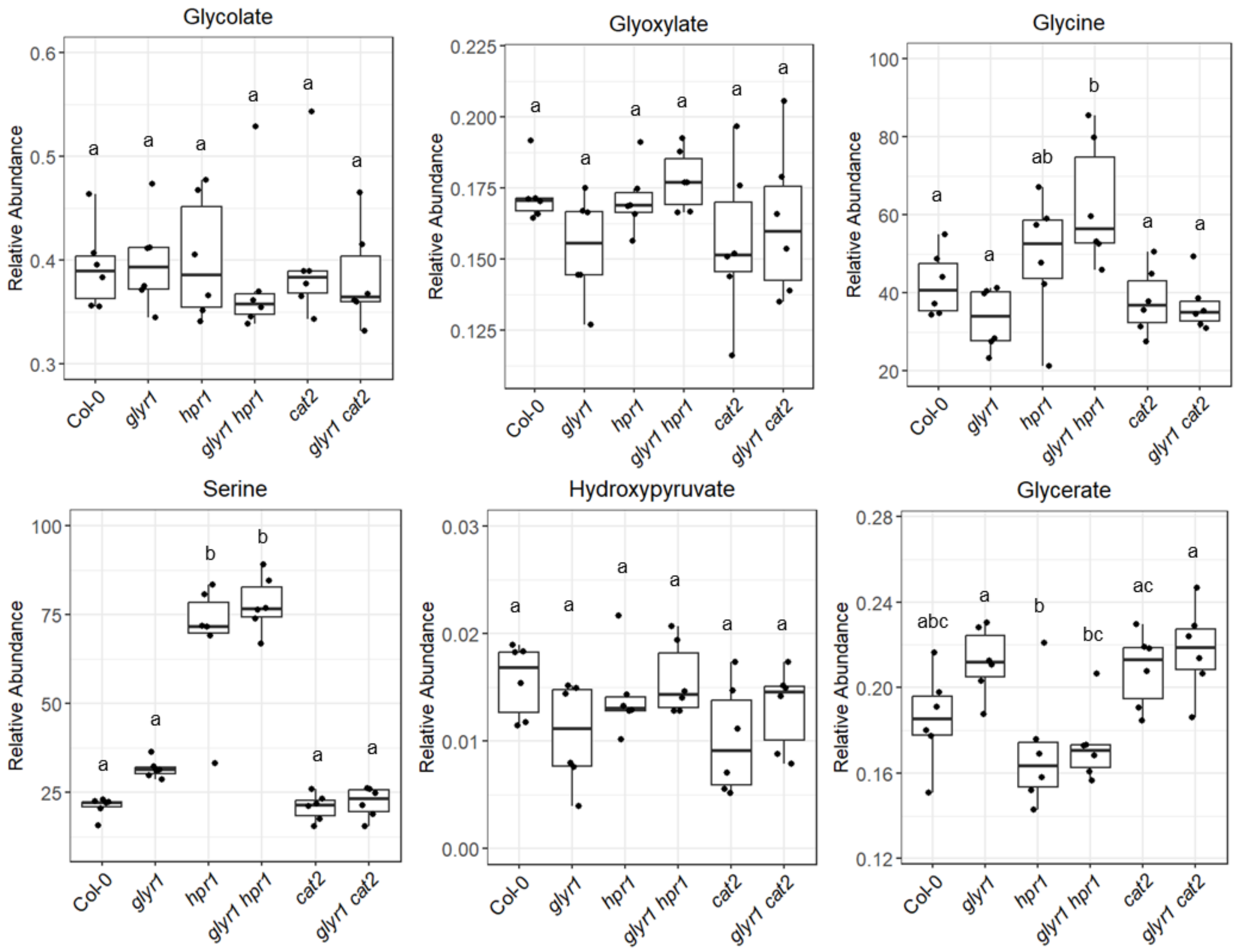


**Supplementary Fig 5. Profiling of photorespiratory metabolites in plants grown under high $CO_2$.** Plants were grown under high $CO_2$ for 3 weeks. Different letters indicate statistically significant differences ($p<0.05$), which were determined by One-way ANOVA with Tukey's HSD test. Biological replicates: n=6.

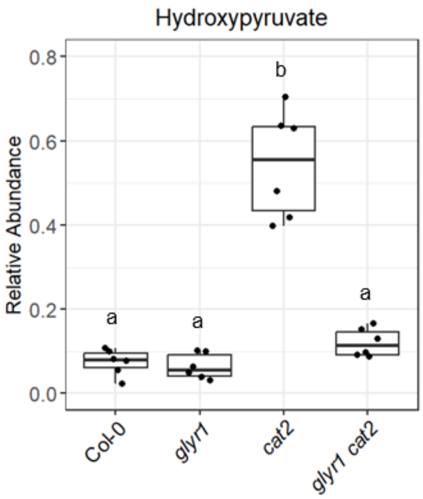


**Supplementary Fig. 6. Profiling of transitional hydroxypyruvate in plants transferred to the photorespiratory environment.** The hydroxypyruvate data used here are the same as those for Fig. 6, except that *hpr1* and *glyr1 hpr1* are removed to focus on *cat2* and *glyr1 cat2*. Plants were grown under 3 weeks of high $CO_2$ and normal light, and then transferred to ambient $CO_2$ and high light before lights were turned on. Leaf tissue was sampled after ~10 h. Different letters indicate statistically significant differences ($p<0.05$), which were determined by One-way ANOVA with Tukey's HSD test. Biological replicates: n=6.

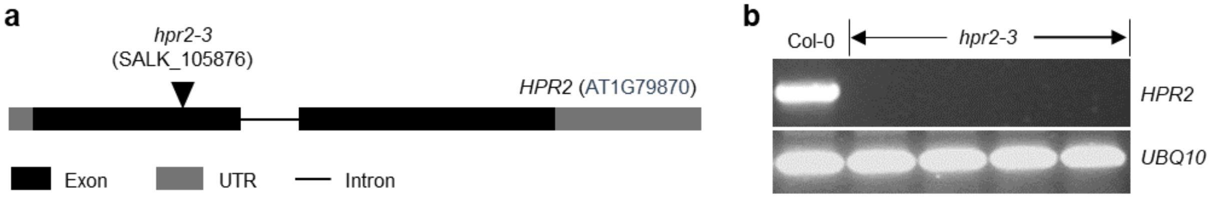


**Supplementary Fig. 7. Characterizations of *hpr2-3*. (a)** Schematic depiction of the *HPR2* gene and the position of the T-DNA insertion in *hpr2-3*. **(b)** *HPR2* expression in plants grown under 2-week normal light followed by 2-week high light. *UBQ10* was used as a control.

**Supplementary Table 1. List of primers used in this study.**

| Primer name | Primer sequence (5'-3') | Purpose |
|---|---|---|
| SALK_LBb1.3 | ATTTTGCCGATTTCGGAAC | Genotyping |
| SALK_067724_LP | GTTGAGTTTGGATATGGCCAC | Genotyping |
| SALK_067724_RP | ACCAAACATCGCGATTACAAC | Genotyping |
| SALK_076998_LP | ACATTTTGGAGCATTGACTGG | Genotyping |
| SALK_076998_RP | TCTGGTGCTCCTGTATGGAAC | Genotyping |
| SALK_053469_RP | GTTTTGCCATAGGCTCGGCTT | Genotyping |
| SALK_053469_LP | CGTCGTCGTCTCCATACCCAT | Genotyping |
| SALK_057410_LP | ACAATCAAAACCCAAAATCCC | Genotyping |
| SALK_057410_RP | AAACGATCTCTTCCCCAAGAC | Genotyping |
| SALK_202680_LP | CTCAGCCAATCCAAATGAGTG | Genotyping |
| SALK_202680_RP | CGGTGTTTTGGAGCAGATATG | Genotyping |
| SALK_203580_LP | GCTTGCAAAAGTTTGATCACC | Genotyping |
| SALK_203580_RP | GTTTGGGAATCATGGGAAAAG | Genotyping |
| SALK_105876_LP | CACTGGATTCCCTAAACATGC | Genotyping |
| SALK_105876_RP | CCCTTAGCTCCTAATGCATCC | Genotyping |
| GLYR1-att-F | ggggacaagtttgtacaaaaaagcaggcttcATGGAAGTAGGGTTTCTGGGT | Cloning |
| GLYR1-att-R | ggggaccactttgtacaagaaagctgggtcCTATTCGCGGGAGAATTTC | Cloning |
| GLYR1-CDS-F | ATGGAAGTAGGGTTTCTGGGT | RT-PCR |
| GLYR1-CDS-R | CTATTCGCGGGAGAATTTCAC | RT-PCR |
| FLAG-GLYR1-F | GACTACAAAGACGATGACGACAAA | RT-PCR |
| UBQ10-F | TCAATTCTCTCTACCGTGATCAAGATG | RT-PCR |
| UBQ10-R | GGTGTCAGAACTCTCCACCTCAAGAG | RT-PCR |
| HPR2-RT-F | ATGGAATCAATCGGAGTCCTTATGA | RT-PCR |
| HPR2-RT-R | CCAAATCCCAAATGTGTCACATGAC | RT-PCR |
